# Genetic variation in correlated regulatory region of Immunity

**DOI:** 10.1101/2022.07.21.500922

**Authors:** Diana Avalos, Guillaume Rey, Diogo M. Ribeiro, Anna Ramisch, Emmanouil T. Dermitzakis, Olivier Delaneau

## Abstract

Studying the interplay between genetic variation, epigenetic changes and regulation of gene expression in immune cells is important to understand the modification of cellular states in various conditions, including immune diseases. Here, we built cis maps of regulatory regions with coordinated activity – Cis Regulatory Domains (CRDs) – in neutrophils, monocytes and T cells. For this, we leveraged (i) whole-genome sequencing (WGS), (ii) chromatin immunoprecipitation sequencing (ChIP-seq), (iii) DNA methylation (450k arrays), and (iv) transcriptional profiles (RNA-seq) from the BLUEPRINT consortium, for up to 200 individuals.

Our study uncovers 9287, 7666 and 5480 histone CRDs (hCRDs) and 6053, 6112, 5701 methyl CRDs (mCRDs) in monocytes, neutrophils and T-cells, respectively. We discovered 15294 hCRD-gene and 6185 mCRD-gene associations (5% FDR). Only 33% of hCRD-gene associations and 37% of mCRD-gene associations were shared between cell-types, revealing the dynamic nature of regulatory interactions and how similarly located regulatory regions modulate the activity of different genes on different cell types. We mapped Quantitative Trait Loci associated with CRD activity (CRD-QTLs) and found that 89% and 70% of these hCRDs and mCRDs are under genetic control highlighting the importance of genetic variation to study the coordination of cellular regulatory programs. We found CRD-QTLs to be enriched in celltype-specific transcription factor binding sites, such as SPI1, STAT3, RFX1, SOX4, ATF3 for neutrophils and monocytes and TCF4 and BCL11A for T-cells, in line with the Human protein Atlas.

We integrated PCHi-C data, which showed that most significant associations discovered within gene-CRD associations and co-expressed genes associated with the same CRD, involving large genomic distances, tend to happen between genomic regions in close spatial proximity. Finally, we mapped trans regulatory associations between CRDs, which enabled the discovery of 207 trans-eQTLs across cell types. Overlapping our hits with trans eQTLs from eQTLGen Consortium meta-analysis in whole blood revealed 52 trans-eQTLs shared between the two studies. Overall, we show that mapping functional regulatory units using population genomics data allows discovering important mechanisms in the regulation of gene expression in immune cells and gain a greater understanding of cell-type specific regulatory mechanisms of immunity.

## Introduction

Genome-wide association studies (GWAS) have identified a large number of genetic variants – mostly located in the non-coding regions of the genome – that are associated with common diseases and complex traits [1]. In addition, extensive collections of genetic variants affecting the transcriptome (i.e. expression quantitative trait loci, eQTLs) across many cell types and conditions are now accessible [2, 3, 4]. Several studies [4, 5, 6] have established the biological mechanisms of eQTLs, describing how non-coding genetic variants alter the activity of regulatory elements, such as through modifications in transcription factor regulation. All these changes present in cis, also propagate along the genome through chromatin interactions, which bring distal elements in close physical proximity [7, 8, 9, 10]. However, there is still much to learn about how genetic variants modulate the regulatory machinery of the cells. Dysregulation of immune and inflammatory activity is present in many complex human diseases. Therefore, analyzing the biological processes of immunity will enable us to understand disease biology and etiology, guiding therapeutic progress. This led us to study the three main primary blood cells of the immune system: neutrophils (key actors of the innate and inflammatory response system), monocytes (which can differentiate into macrophages and dendritic cells to trigger an immune response) and T-cells (essential part of the adaptive immune system).

To understand gene expression regulatory machinery and eQTLs upstream mechanisms, studies have analyzed the inter-individual variation of histone modifications, using ChIP-seq data [11, 12] to discover the existence of coordinated activity of sets of regulatory elements, called Cis-Regulatory Domains (CRDs). Others have leveraged population variation of chromatin accessibility using ATAC-seq libraries [13, 14, 15, 16], as it explains 70% of gene expression variance. Since DNA methylation is thought to influence chromatin structure [17] and gene expression when located in regulatory regions [18], the study of DNA methylation variability across a population [19] also provides tools to infer the mechanisms underlying transcriptional regulation. In addition, many studies highlighted the role of transcription factors (TFs) in regulating gene expression [20].

In order to gain further insight into the molecular processes at play, and shed light on the cell-specificity of such regulatory mechanisms, we leveraged multiple omics data from the BLUEPRINT Consortium [21], for the three key primary immune cell-types mentioned above. This dataset includes whole-genome sequencing (WGS), chromatin immunoprecipitation sequencing (ChIP-seq) for two histone modification marks associated with active enhancers and promoters (H3K4me1 and H3K27ac), DNA methylation (Illumina 450K arrays) as well as transcriptional profiles (RNA-sequencing) for more than 197 individuals. In addition, we exploit promoter capture Hi-C (PCHiC) datasets for the same cell types [22]. This extensive dataset enables the study of population-wide perturbations which modifies the transcriptomic dynamics at play.

In this paper, we mapped CRDs in 3 primary immune cell lines, to investigate the cell-type specificity of such regulatory structures and their role in the modulation of gene expression. We discovered a large amount of CRD-gene associations revealing the dynamic nature of regulatory interactions and how similarly located regulatory regions modulate the activity of different genes on different cell types. Additionally, we highlighted the role of genetic variation in the coordination of cellular regulatory programs, and consolidated the functional interactions discovered by integrating PCHi-C and showing that these associations take place in close physical proximity. Finally, we leveraged the trans-CRD networks and TRHs discovered to infer inter-chromosomal interactions and show that this could be used to discover trans-eQTLs. Overall, we show that mapping functional regulatory units using population genomics data allows discovering important mechanisms in the regulation of gene expression in immune cells.

## Results

### A map of cis-regulatory domains in 3 primary immune cell types

We subsampled histone peaks from 250 ChIP-seq assays (for three histone modifications and 3 cell types) (see Methods), to create a consensus set of peak coordinates. Then, to define the map of cis-regulatory domains (CRDs) in primary immune cells, we used a previously published method [12], which relies on the hierarchical clustering of molecular phenotype data across a population of individuals, and creates groups of peaks exhibiting high correlation (see Methods). The subsequent tree was cut into areas of high correlation and covering at least two different chromatin regions. Applying this framework to histone ChIP-Seq data, we discovered 9287, 7666, 5701 histone CRDs (hCRDs) in monocytes, neutrophils and T-cells, respectively [Figure 1 and Supplementary 1A, 1B]. Since the histone marks (H3K4me1 and H3K27ac) are associated with active enhancers and promoters, these histone CRDs capture the correlated activity of these regulatory elements. We extended this method to another type of epigenomic data: DNA methylation, obtained through Illumina 450K arrays, and found 6053, 6112, 5701 methyl CRDs (mCRDs) in monocytes, neutrophils, and T-cells, respectively [Supplementary Table 1A]. As CpG islands are preferentially located within promoter regions, mCRDs correspond to the synchronized activity of promoters.

**Figure 1:**
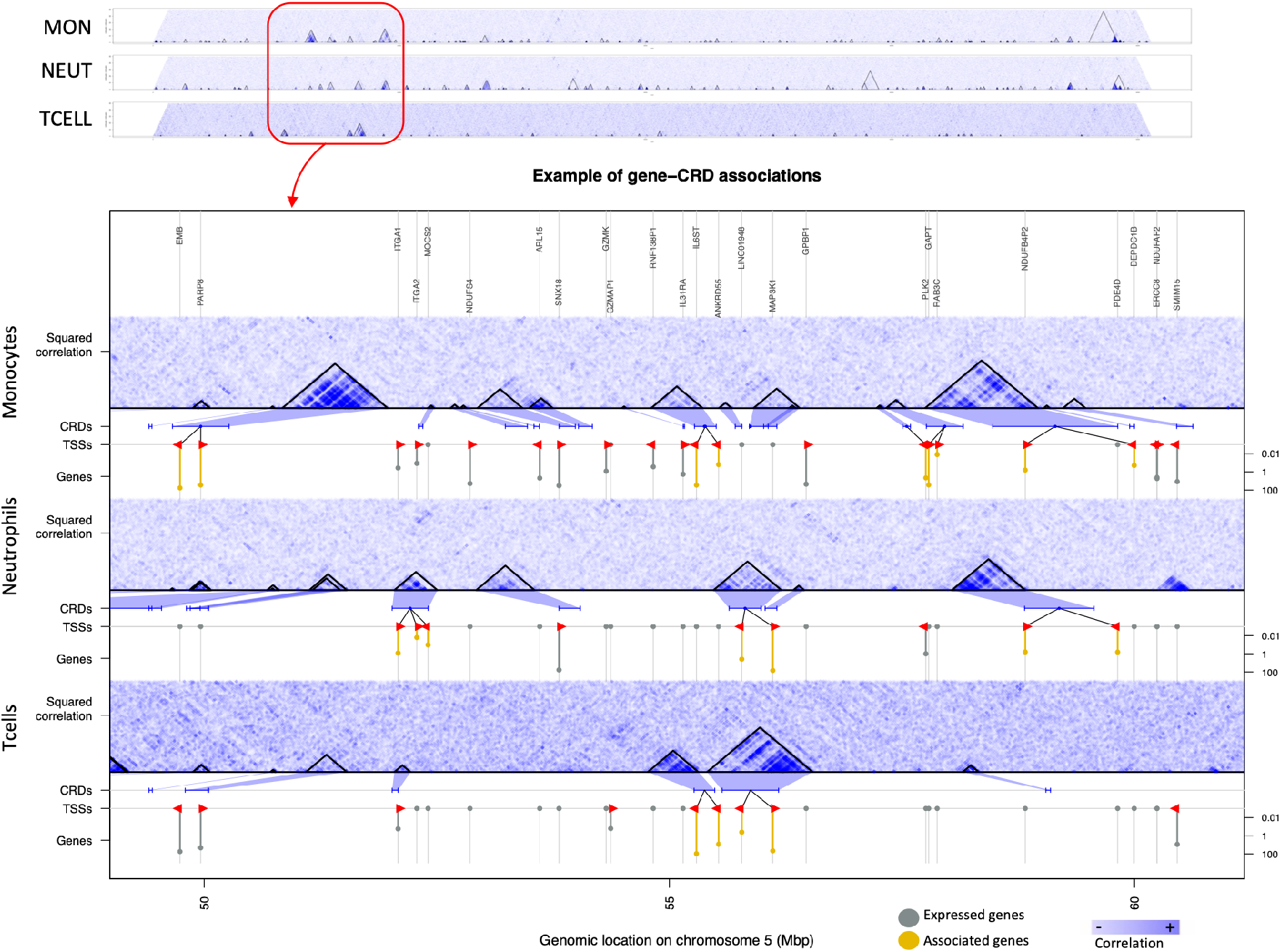
hCRDs emerging from the inter-individual correlation between chromatin peaks for Monocytes, Neutrophils and T-cells. CRDs represent the coordinated activity between nearby regulatory elements (promoters and enhancers). Chromosome 5 is represented here, with a zoom of a region spanning 1000 chromatin peaks. hCRDs are outlined by black triangles, and the hematopoietic lineage is represented on the right. Significant gene-CRD associations (5% FDR) are represented in red. For each celltype, expressed genes are colored in grey and significantly associated genes are colored in yellow.

As expected, the different resolution of DNA methylation data compared to histone ChIPSeq (single base pairs vs kilobases) affects the size distribution of CRDs. We found that hCRDs exhibited a unimodal distribution centered around 40 kb, whereas mCRDs displayed a bi-modal distribution, peaking at 300bp and 40kb [Supplementary Figure 1C]. To correct for this, we modeled the distribution using a mixture gaussians for small (0.2 to a few kb) and large (a few kb to 1-2Mb) domains. As CRDs involve at least two distinct, non-overlapping regulatory regions [12], we discarded the mode centered on 300bp likely representing CpG sites located in the same regulatory region. Therefore, we chose a threshold corresponding to the 0.95 percentile of the distribution of small domains, to define a set of methyl CRDs we selected for downstream analysis.

Given that sample size could be an important factor in the discovery of CRDs, we subsampled our dataset to the lowest sample size available (N=94) for the discovery of CRDs, and also integrated the SySGenetix LCL dataset [11] as a baseline. In agreement with previous analysis showing that about 100 individuals are sufficient to map more than 90% of CRDs in LCLs [12], we found that the difference in the number of CRDs discovered across cell types was maintained before and after subsampling for hCRDs (89% of hCRDs discovered with 94 samples) but less for mCRDs (67% of mCRDs discovered with 94 samples) [Supplementary Table 1A].

To investigate the patterns of sharing of CRDs between cell types, we labeled a CRD as shared between two cell types if at least 50% of its histone peaks (hCRDs) or CpG islands (mCRDs) in the query cell-type were also present within a CRD in the reference cell-type. We integrated the SysGenetiX LCL dataset to our analysis (subsampled to 94 samples) and analyzed CRD sharing in a pairwise fashion for all cell-types [Supplementary Table 1A, Supplementary Figures 1D,E]. We found that T-cells and LCLs shared the most CRDs, with a mean of 32% among hCRDs, which was expected as they both descend from a common lymphoid progenitor. Similarly, neutrophils and monocytes shared 24% of their CRDs, and these cells descend from a common myeloid progenitor. These results are therefore consistent with the hematopoietic lineage. Of note, we confirmed these results by analysing individual epigenomic marks within CRDs (histone peaks or CpG sites) instead of CRDs and by measuring their overlap between cell-types [Supplementary Figures 1F,G]. Indeed, neutrophils and monocytes are both formed from myeloblasts and are part of the innate immune system whereas T-cells and LCLs (immortalized cell-lines descending from B-cells) belong to the adaptive immune system. In addition, LCLs hCRDs displayed an important overlap with primary immune cells (24, 27 and 32% in neutrophils, monocytes and T-cells), due to the number of CRDs discovered, substantially more important than any other cell type (10497, 5480, 7660,6831 hCRDs discovered in LCLs, T-cells, monocytes and neutrophils, respectively, when subsampling the dataset to the lowest sample size across cell types (N=94), or 12583, 5480, 9287 and 7666 otherwise) [Supplementary Table 1A]. Notably, we reached similar results in terms of lineage and overlap for mCRD sharing among cell-types, with 34% of mCRDs shared between neutrophils and monocytes, and 18 to 25% of mCRD sharing between T-cells and the other cell-types. Overall, mCRDs displayed a greater amount of tissue sharing than hCRDs, indicating that they may be less tissue specific than the histone CRDs [Supplementary Figure 1E]. This is consistent with the previous finding that promoter activity is less cell type-specific than enhancer activity [23]. Overall, our results demonstrate that hCRDs are highly cell-specific as they capture coordinated activity of enhancers and promoters, mCRD are less cell-specific as they only capture this for promoters, and the amount of sharing between cells recapitulates the hematopoietic lineage.

### Regulatory interaction dynamics across immune cell types

CRDs represent areas of high correlation between epigenomic marks within a population. We computed hCRD activity, by averaging the activity of the chromatin peaks within an hCRD per individual, and therefore obtaining a quantification vector per CRD. We applied a similar process to CpG sites within mCRDs to obtain their respective activity. To investigate the role of CRDs in the regulation of gene expression in immune cells, we correlated the CRD activity with gene expression. We performed the analysis in cis, considering genes within 1Mb of the CRD in each cell-type to investigate their role in the regulation of gene expression. We discovered a large amount of CRD-gene associations, with respectively 6300, 6755, 2239 hCRD-gene associations and 2027, 2300, 1858 mCRD-gene associations in neutrophils, monocytes and T cells at 5% FDR [Figures 1, 2].

**Figure 2:**
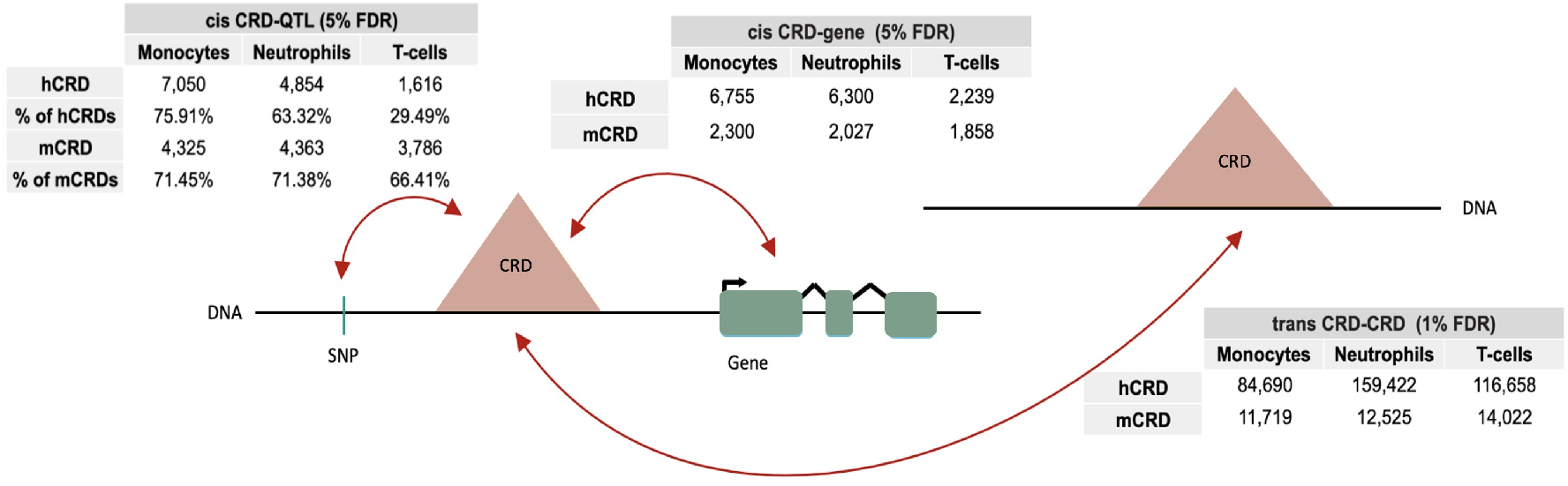
Schematic of CRD associations (CRD-QTLs, CRD-gene and CRD-CRD) with the number of associations found within the maximum sample size available for each cell type and epigenomic mark.

Next, we investigated the cell specificity of the regulatory machinery on the transcriptome. To compare CRD-gene associations between two cell types, we quantified CRDs cell-type specific activities by considering the activity of the same exact set of peaks across the three cell types. The fraction of significant CRD-gene associations in common between the 2 cell types was then calculated over the total number of significant CRD-gene associations of the reference cell (FDR 5%). We found that neutrophils and monocytes shared between 33% and 37% of their hCRD-gene and mCRD-gene associations [Supplementary Figures 2A,B]. Conversely, the fractions of significant CRD-gene associations of monocytes and neutrophils with T-cells were smaller (from 10 to 31%), again consistent with the hematopoietic lineage. Furthermore, these findings reveal the dynamic nature of regulatory interactions across cell-types, and how similarly located regulatory regions modulate the activity of different genes on different cell types.

As a previous study [23] showed that chromatin modification profiles in enhancers are highly cell-type specific whereas most genes expressed are shared between these cells, we compare the cell-specificity of CRD-gene associations and genes expressed between cell-types. We extracted the genes significantly associated with CRDs in each cell type, and looked at the intersection of these gene sets. We found up to 47% of genes in these gene-CRD associations were shared with at least one other cell type [Supplementary Figures 2A,B]. As less CRD-gene associations than genes are shared between cell-types, this indicates that gene-CRD associations are more tissuespecific than the sets of genes expressed between cell types in those associations, highlighting the reprogramming of gene regulation through chromatin organization in the hematopoietic lineage.

We subsequently investigated the complexity of connectivity between genes and CRDs across cell types and found that 32.1% of hCRDs associated with at least one gene in a given cell type, and 13.7% of hCRDs associated with two or more genes [Supplementary Figures 2C]. Conversely, we found 23.3% of genes associated with at least one hCRDs and up to 6% with at least two hCRDs. Similarly, we found around 16.2% of mCRD associated with at least one gene, and 9.4% of genes associated with at least one mCRD. These results highlight the complexity of regulatory relationships among genes and CRDs.

As Ribeiro et al. [24] integrated ChIP-seq and Hi-C to interpret the molecular mechanisms driving gene co-expression, we further characterized the impact of CRDs on the complexity of gene regulation, and aimed to quantify how much of gene co-expression is driven by shared CRD regulation. We looked into the 29940, 46146 and 13737 cis co-expressed gene pairs we found (i.e. genes whose expression is correlated among individuals) (FDR 1%) respectively for neutrophils, monocytes and T-cells [Supplementary Table 2D]. We then asked how their position relative to associated CRDs affected their pattern of association and we found that the fraction of co-expressed genes associated with the same CRD was strongly enriched at long distances(100 to 1000kb) [Supplementary Figure 2E], suggesting that interaction with a given CRD is an important mechanism for the distal coordination of gene expression. Furthermore, we notice that most genes are located within the CRD they are regulated by. Gene pairs odds ratios of belonging to the same CRD while being co-expressed are quite high (ranging from 11 to 286), comforting the fact that CRDs are major regulators of co-expression.

### CRDs are under genetic control

To investigate how genetic variation affects the regulatory machinery, we tested whether CRDs are under genetic control. For this, we mapped CRD-QTLs finding that around 60% of hCRDs and 70% of mCRDs were genetically controlled with respect to their overall activity [Figure 2]. Overall, mCRDs are more genetically controlled than hCRDs, which could be explained by the proximity of methylation CpG marks to promoter regions, where the genetic signal may be stronger. Furthermore, CpG marks are obtained through Illumina arrays which only sample selected areas for methylation marks. The patterns of CRD-QTLs sharing among tissues were also estimated using the *π*_1_ estimate (the proportion of true positives [25]), extracting the significant CRD-QTLs identified in one cell-type and then tested for replication in another cell type (median hist *π*_1_=0.66, median methyl *π*_1_=0.51) [Supplementary Figure 2F]. These results reveal that most genetic variants controlling the activity of the CRDs are shared between cell-types. We wondered whether cell-type specific CRD-QTLs were controlled by cell-type specific transcription factors (TFs). We incorporated knowledge from external databases such as Motifmap [26] and Remap [27], and extracted the 50 TFBSs that overlapped the most with ChIP-seq peaks located within CRDs. Computing enrichment for the significant CRD-QTLs found within these 50 most represented TFs, and subsequently calculating Fisher’s test odds ratios, we found significant enrichment for 18 different TFBSs (FDR 1%) [Supplementary Figures 2G,H]. We found that monocytes and neutrophils CRD-QTLs were enriched in the transcription factor binding sites of SPI1, STAT3, RFX1, SOX4, ATF3, with odd ratios higher than for T-cells. All of these TFs are enriched in monocytes and neutrophils, according to the Human Protein Atlas [28]. Conversely, TCF4 and BCL11A elicited stronger odd ratios within T-cells and monocytes, again in line with the Human Protein Atlas which labels these TFs as group-enriched in lymphocytes and dendritic cells (descending from monocytes in the hematopoietic lineage). These findings suggest that CRD activity changes are driven by genetic modification in transcription factor binding sites.

### CRD structure and connectivity reflect functional 3D chromatin organization

As our previous work [12] indicated that CRDs are associated with functional 3D interactions, we investigated how CRDs in primary immune cells are related to 3D genome structure using promoter capture Hi-C data from Javierre et al., Cell 2016 [22].

CHiCAGO algorithms perform normalization and multiple testing specifically adapted to CHi-C experiments [29] and consider PCHi-C interactions significant if the interactions detected by CHiCAGO have a score superior or equal to 5. We found that the fraction of correlated chromatin peak pairs (per chromosome pair-wise association testing, FDR 1%) increased with the PCHi-C interaction score (PCHi-C interactions are considered significant if the CHiCAGO score≥ 5) [29] [Supplementary Figure 3A]. Moreover, correlated histone peak pairs were more likely to be in close 3D physical proximity (CHiCAGO score ≥ 5) than uncorrelated peaks, with the effect maximizing for pairs separated by 50-500kb [Figure 3A and Supplementary 3B,C] (for distances ¡20kb, we expect genomic distance noise to affect short-range PCHi-C interactions [30], therefore we do not consider this distance interval in the analysis).

**Figure 3:**
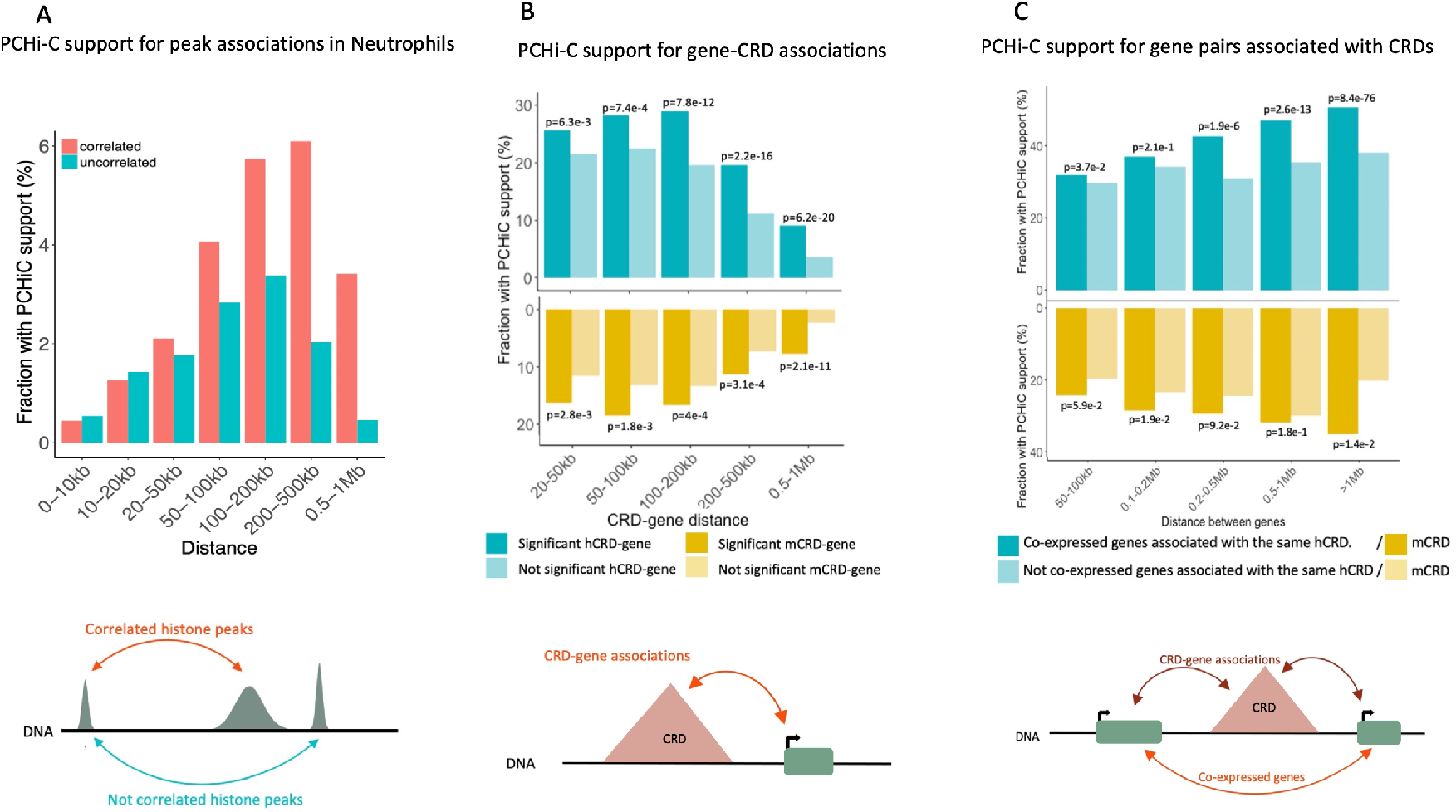
A. Fraction of neutrophil chromatin peak pairs on the same chromosome supported by PCHi-C data (CHiCAGO score ≥ 5) at significantly associated (pink) and non-associated (blue) pairs of chromatin peaks within bins of increasing distance between peaks. B. Fraction of hCRD-gene and mCRD-gene associations supported by PCHi-C data (CHiCAGO score ≥ 5) at increasing CRD-gene distances. C. Fraction of of hCRD-gene associations and mCRD-gene associations supported by PCHi-C data (mean CHiCAGO score ≥ 5) for pairs of co-expressed genes (5%FDR) that associate with the same CRD. The fraction is measured at bins of increasing distance between co-expressed genes.

Then, we explored how cis gene-CRD associations may reflect long-range physical interactions between regulatory elements and gene promoters. We observed around 30%-40% of gene-CRD associations were supported by PCHi-C (CHiCAGO score≥ 5), indicating a strong enrichment compared to correlated peak pairs [Figure 3B]. In addition, we notice that mCRD-gene associations are less supported by PCHi-C than hCRD-gene associations (around 15% less PCHi-C support). We observe an increase in PCHi-C support when gene-CRD distance is between 50-200kb, especially for T-cells associations, indicating that gene-CRD associations at large genomic distances tend to happen between genomic regions in close spatial proximity.

As shown before, nearby genes are often co-expressed, and these associations have been mapped across 49 human tissues [24]. To get a better understanding of these molecular mechanisms, we mapped pairs of co-expressed genes associated with the same CRD and we found that these associations showed increasing PCHi-C support with increasing genomic distance between genes [Figure 3C]. For genes inside a CRD, 20% and 36% of co-expressed gene associated to a hCRD and mCRD were supported by PCHi-C. For distances between 50kb and 1Mb between co-expressed gene pairs, the fraction of pairs supported by PCHi-C increases, supporting once more that associations at large genomics distance is therefore compensated by spatial proximity.

### Trans CRD networks highlight cell-type specific biological functions in immune cells

Given that regulation of gene expression in trans is an important component to the overall variance in gene expression [31], we mapped CRD trans networks in immune cells.

We first computed inter-chromosomal pairwise CRD trans associations in each cell type to discover respectively 159’422, 84’690 and 116’658 significant hCRD-hCRD trans-associations (FDR 1%) and 12’525, 11’719 and 14’022 significant mCRD-mCRD trans-associations (FDR 1%) in neutrophils, monocytes and T cells [Figure 2]. We repeated the same analysis using a fixed sample size (n=94) for all cell types and found that even though the number of significant associations depended strongly on sample size, important differences between cell types were maintained [Supplementary Figure 3D].

We then built trans CRD networks using the discovered associations as edges between CRDs (nodes) [32] to define Trans Regulatory Hubs (i.e. communities) within those networks. Using this approach, we found between 31 to 308 TRHs in hCRDs, and over 200 TRH per cell type in mCRDs, depending on the cell type [Sup. Table 1A, Supplementary Figure 3E]. We further observed that each cell type had a few TRHs with more than 1000 CRDs [Supplementary Figure 3F], while most of the individual hCRDs had only a small number of associations each to other CRDs [Supplementary Figure 3G] indicating regulatory networks of very large complexity.The patterns of trans CRD-CRD association sharing were also estimated by extracting the significant trans CRD associations identified in one cell-type and then tested for replication in another cell type [Supplementary 3H,3I]. (median *π*_1_=0.35 for trans hCRD sharing, and 0.92 for trans mCRD sharing). The subsequent results showing 35% of trans hCRD-hCRD sharing between cell types comfort our findings of highly cell-type specific maps of gene regulation.

Since we established that trans CRD associations identify functional chromatin interactions and CRDs coordinate distal gene expression, we investigated whether genes belonging to the same TRH have similar biological properties, i.e. participate in the same biological processes, perform similar molecular functions, or their protein products colocalize in the same cell region. To this end, we researched if some TRHs showed an enrichment in immune processes. We performed gene set enrichment analysis of the genes associated with TRHs of hCRDs, using the GOrilla algorithm [33] and REVIGO [34] visualization platform to determine over-represented gene categories. We defined as background gene sets, all the genes expressed for each cell type, and the target gene sets were defined as all the genes significantly associated with the CRDs of the TRH studied. Out of 122 TRHs with genes associations (84 for neutrophils, 6 for macrophages, 22 for T-cells), we selected the 25 largest TRHs from which 10 showed significant gene ontology (GO) term enrichment at 1% FDR (13 TRHs at 5% FDR). We discovered 2 TRHs, one in neutrophils and one in T-cells, which are linked to immune response associated GO terms (31 terms at 1% FDR and 64 at 5% FDR) [Supplementary 3J,3K]. These include the regulation of T-cell differentiation, lymphocyte differentiation, regulation of adaptive immune response, and regulation of leukocyte cell-cell adhesion. The strongest signal was associated with the regulation of adaptive immune response in T-cells (29 GO terms at 1%FDR). We noticed that the majority (9 out of 13, FDR 5%) of TRHs have their genes performing similar molecular functions, supporting the fact that genes linked to the same TRH participate in correlated biological processes. Specifically, we observed a T-cell TRH (index 3, 421 genes) consisting of genes involved in the regulation of cell communication, regulation of cell-cell adhesion, cell surface receptor signaling pathway, and colocalization of gene products in the plasma membrane and cell surface. These findings underscore the cell-cell signaling pathway in T-cells as the underlying biological mechanistic process. Once the T-cell receptors bind an antigen, the cell will activate a series of internal signaling pathways that allow for the antigen recognition to be verified, leading to the proliferation of T-cells specific to this antigen [35]. These results show that trans CRD networks and TRHs are key factors in the regulation of gene expression and provide a mechanistic explanation for biological pathways.

### Mapping trans eQTLs through histone Trans CRD networks unravel new trans eQTLs

We previously showed that integration of trans CRD-CRD associations with eQTL analysis and CRD-gene associations could successfully indicate candidate trans SNP-gene pairs with higher prior probability of being positive in trans-eQTL mapping [12]. Expanding on this approach, we studied the propagation of trans-eQTL effects in three primary immune cells, and selected the two most likely trans-eQTL causal models [Supplementary 3L] out of the 18 tested ones in the study of Delaneau et al., Science 2019 [12]. The first scenario consists of a QTL variant associated with a CRD in cis, which is associated to another CRD in trans, and the latter CRD is associated to a gene in cis. The second scenario involves a cis association between the gene of a cis-eQTL and a CRD, which in turn is trans associated to another CRD cis-asscoiated to a gene. Out of 265k, 176k and 46k tested associations of scenario1 in neutrophils, monocytes and T-cells we found 55,16 and 5 significant associations (5% FDR). We found an enrichment of small p-values for the combinations we evaluated. In particular, we found that neutrophils had the largest number of hits with 55 and 62 trans-eQTLs using respectively an hCRD-QTL (scenario 1) and an eGene (scenario 2) [Supplementary 3M].

Overlapping our hits with the trans eQTLs from a meta-analysis in whole blood (eQTLGen Consortium) [37] revealed an important number of signals shared between the two studies. Indeed, we found 171, 81 and 9 unique trans-eQTLs in neutrophils, monocytes and T-cells. A third of the trans-eQTLs discovered in neutrophils overlapped with the eQTLGen trans-eQTLs, while monocytes and T-cells had only one association in common, involving the same gene and a variant in linkage disequilibrium (LD) with our variant (LD¿0.5) [Supplementary 3N]. As neutrophils are the most abundant cell type in whole blood after red blood cells, they might be prone to contributing the most to whole blood gene expression patterns [36]. These results therefore indicate that our data integration strategy is able to discover trans-eQTLs overlapping with existing datasets.

## Discussion

Understanding the regulation of gene expression in human cells is key to mechanistically deciphering how gene expression patterns arise and lead to cell type identity in health and disease. In particular, genetic variation in regulatory elements offers the possibility to study populationwide perturbations to those gene networks and infer their global structure and patterns. In a study from 2016, Chen et al. [4] analyzed the same three immune cells from the Blueprint dataset to investigate the genetic and epigenetic effects on RNA transcription and splicing. Here, we explore the regulation of the transcriptome at another scale, and emphasize the cell-specificity of the biological mechanisms at play. Furthermore, extending the CRD framework [12] to methylation data enables the discovery of mCRD networks, which facilitates the analysis of large datasets in which histone modifications are not available. In this context, mapping of CRDs and TRHs from epigenomics data (histone modifications and DNA methylation marks) enable the partition of the genome in a given cell type in cis and trans functional regulatory units.

Applying this framework to immune cells revealed that those structures are present in high numbers and show high tissue specificity. Furthermore, the cell-specificity of the observed dynamics are in accordance with the hematopoietic lineage, which displays more resemblance between neutrophils and monocytes than with T-cells, as they both derive from myeloid progenitor.

In particular, we detected a large number of trans-associations and TRHs, which allowed inferring inter-chromosomal interactions and discovering cell-type specific trans-eQTLs. We observed that almost a third of the trans-eQTLs discovered in neutrophils overlapped with the eQTLGen trans-eQTLs, while monocytes and T cells had only one association in common. This result is consistent with neutrophils being the most abundant cell type in whole blood (excluding red blood cells), thereby likely contributing the most to whole blood gene expression patterns.

Association of CRD activity with expression of genes in cis revealed that CRDs play an important role in tissue-specific gene expression, with an effect especially strong at long distances. The observation that, in neutrophils, the majority of co-expressed genes separated by 100kb to 1Mb are linked to the same CRD is an important example of the functional role of CRDs in gene expression. Moreover, the fact that a substantial fraction of CRD-gene associations is involved in 3D contacts supports the hypothesis that CRDs reflect functional regulatory interactions in the 3D genome.

Taken together, our results highlight the power of generating cell-type specific maps of regions of coordinated activity to discover important biological mechanisms. It indeed reduces the search space by several orders of magnitude, so that the discovery of relevant interactions becomes accessible to standard statistical association testing. We therefore surmise that the use of population genomics data will be key to define functional regulatory units in a tissue-specific manner and will allow for more relevant and targeted analysis of the regulatory potential of the human genome. In this respect, the methodology presented in this paper could be applied to the widely available population DNA methylation data in order to rapidly create CRD and TRH maps in a large number of tissues and conditions. Although our approach is strictly computational and further experimental validation is required to link non-coding variants and regulatory elements to genes involved in the dysregulation of immune processes, we provide a way to improve our understanding of the cell-specific mechanisms of the regulatory machinery, to reveal the disrupted biological mechanisms in disease and pave the way for new therapies.

## Acknowledgements

Funding: This work was supported by grants from SYSCID (a systems Medicine approach to chronic inflammatory Diseases) under the European Union’s Horizon 2020 research and innovation program (grant agreement no. 733100). The founders had no role in study design, data collection and analysis, decision to publish, or preparation of the manuscript.

## Author contributions

E.T.D. and O.D. designed and supervised the study and contributed to data interpretation. D.A. and G.R. designed and executed the primary data analysis. A.R. contributed to data analysis. D.A., G.R., D.M.R., O.D and E.T.D. performed the primary manuscript writing.

## Methods

### Sample collection

Genotyping, DNA methylation, RNA-Seq, and two ChIP-seq experiments, H3K4me1 and H3K27ac, have been performed in neutrophils, monocytes and T cells in the scope of the Blueprint Epigenome Consortium. The data have been downloaded from the European Genome-Phenome Archive.

We used the LCL data from from the SysGenetix consortium [12].

We integrated PCHi-C data from Javierre et al., Cell 2016 [22]. Their PCHi-C libraries were sequenced on the Illumina HiSeq2500 platform, then reads were processed using the HiCUP pipeline [38], which maps the positions of di-tags against the human genome (GRCh37), filters out experimental artifacts, such as circularized reads and re-ligations, and removes all duplicate reads. Interaction confidence scores were computed using the CHiCAGO pipeline [29]. Briefly, CHiCAGO calls interactions based on a convolution background model reflecting both ‘Brownian’ (real, but expected interactions) and ‘technical’ (assay and sequencing artifacts) components. The resulting p values are adjusted using a weighted false discovery control procedure that specifically accommodates the fact that increasingly larger numbers of tests are performed at regions where progressively smaller numbers of interactions are expected. The weights were learned based on the decrease of the reproducibility of interaction calls between the individual replicates of macrophage samples with distance. Interaction scores were then computed for each fragment pair as –log-transformed, soft-thresholded, weighted p values. Interactions with a CHiCAGO score ≥ 5 in at least one cell type were considered as high-confidence interactions.

The list of significant trans-eQTLs (FDR ≤ 0.05) from the eQTLGen consortium was downloaded from the eqtlgen.org website.

### Data preparation:ChIP-seq data and Methylation data

One of the requirements of this study is to get a population scale call set of chromatin peaks. This implies that the coordinates of chromatin peaks need to be similar and comparable across samples and cells. We first determined the peak coordinates and then quantified each individual according to the peak coordinates. To build a population call set of peaks, we first build for each ChIP-seq assay (H3K4me1 and H3K4me1) a ‘consensual individual’ by aggregating 1e6 randomly sampled ChIP-seq reads from 50 neutrophils, 50 monocytes and 50 T-cells, together in a unique BAM file (therefore containing 150e6 ChIP-seq reads). Then, we carried out the actual peak calling onto this ‘meta’ BAM file in order to get a consensus set of peaks across multiple individuals and cell types (leading to the identification of 66770 H3K27ac peaks and 91528 H3K4me1 peaks). This particular step has been done using the program findPeaks from the software package HOMER v4.9 (webpage: http://homer.ucsd.edu/homer/ngs/peaks.html), parameterized with options adapted for histone marks: -style histone -o auto. We repeated the procedure for each ChIP-seq assay.

Once all chromatin peaks coordinates are known, we proceeded with the per-sample quantification. To do so, we used the script annotatePeaks.pl from the software package HOMER v4.9 [39] in combination with the peak coordinates we determined. This script counts the number of ChIP-seq reads falling within the peak coordinates. We run this script using the following options: noann -nogene -size given independently for each individual and each ChIP-seq assay and obtained per-peak read counts that were subsequently normalized over the reads per sample/assay pair. We then assembled the data into six quantification matrices: the normalized read counts across the 2 ChIP-seq assays and the three cell types.

For the integrated analysis of the BLUEPRINT and SGX datasets using the same sample size (n=94), we randomly selected 94 individuals for each cell type and perform Molecular phenotype data preparation in a similar way and identified 71009 H3K27ac peaks and 106067 H3K4me1 peaks.

### Data preparation:RNA-seq data

The read mapping of the Blueprint data set was carried out using GRCh37/hg19. Gene expression was quantified from BAM files using Qtltools quan function. We then filtered out genes that were poorly quantified across samples by removing all genes with more than 10% null RPKM values across samples. Finally, we only kept the genes in downstream analyses that are either protein coding or long non-coding RNAs given the GENCODE v15 annotation.

### Covariate correction and normalization of molecular phenotypes

The variability in molecular phenotypes (RNA-seq, ChIP-seq and DNA methylation data) can be from either technical or biological origin. The idea here is to correct for technical variability only while retaining as much as possible of the biological variability. In other words, the goal is to maximize the signal-to-noise ratio. Covariate correction of molecular phenotype data was performed in a similar way to Delaneau et al., Science 2019. Briefly, we residualized the molecular phenotype data for two types of covariates described below: 1. Sex. We used the metadata provided in EGA.

2. Experimental. We performed PCA on each of the quantification matrices independently and used the individual coordinates on the first PCs as covariates. We sequentially corrected molecular phenotype data using 2 to 50 PCs and used the number of PC that maximize the number of QTL discovered.

Finally, quantification matrices were rank-normalized on a per phenotype basis across all individuals so that quantifications match a normal distribution.

We selected 10 PCs to correct our ChIP-seq quantification matrices using QTLtools correct and then merged the ChIP-seq quantifications matrices to have 1 matrix per cell type. We also selected 10 PCs for the expression arrays and 12PCs for the methylation arrays.

### Building correlation and CRD maps

CRD were called with the method developed by Delaneau et al., published in Science in 2019 (https://github.com/odelaneau/clomics) [12].

First, we built correlation maps by measuring interindividual correlation between ChIPseq peaks located in the same chromosome with a 250 peaks sliding window (and retrieving the Pearson correlation coefficients using corrected and rank-normal transformed data matrix). Then, an agglomerative hierarchical clustering algorithm is applied per chromosome, in which ChIP-seq peaks are assigned to clusters and iteratively the clusters are merged as we move up in the hierarchy. This strategy resulted in a binary tree that regrouped all ChIP-seq peaks from the same chromosome in which each node delimited a set of highly correlated ChIP-seq peaks.

CRDs were called by identifying the minimal set of internal nodes that captured most of the overall correlation mass (i.e., cumulative sum of squared correlation). To retain an internal node as a CRD, three criteria needed to be fulfilled: i) CRDs regrouped only highly correlated ChIP-seq peaks: the mean absolute correlation between all possible pairs of ChIP-seq peaks within a CRD had to be at least twice as high as the mean correlation between all ChIP-seq peaks in the chromosome; ii) CRDs had well-defined boundaries: the mean absolute correlation between all pairs of ChIP-seq peaks involving either the first or the last ChIP-seq peak (on the basis of their genomic location) had to be at least twice as high as the same value derived for the first and last peaks on the chromosome; iii) CRDs captured distal coordination between at least two regulatory elements (REs): ChIP-seq peaks had to cover at least two non-overlapping regulatory regions. These three criteria were implemented into an algorithm that processed each binary tree starting from the root node (node regrouping all peaks of a chromosome) and recursively traversed the internal nodes of the tree until an internal node fulfilled all three criteria. Then, declared the internal node and all the peaks downstream as a CRD, stopped to go deeper by ignoring the children of this node and carried on with other internal nodes in the tree.

This pipeline was applied to histone peaks as well to CpG islands. Therefore, we obtained histone CRDs (hCRDs) and methyl CRDs (mCRDs).

We found that hCRDs exhibited a unimodal distribution centered around 40 kb, whereas mCRDs displayed a bi-modal distribution, peaking around 300bp and 40kb. We thus modeled the distribution using a mixture gaussian model of small domains (0.2 to a few kb) and large domains (a few kb to 1-2Mb). As CRDs were defined as domains involving at least two distinct non-overlapping regulatory regions, we discarded the mode of 300bps likely representing CpG sites located in the same regulatory region. We set a threshold corresponding to the 0.95 percentile of the distribution of small domains, to define a set of size-selected methyl CRDs.

### CRD specificity across cell types

For determining hCRD and mCRD sharing between different cells, we compared ChIP-seq peak correlation maps and CpG sites correlation maps between cells. A CRD is shared between 2 cells if 50% of the peaks belonging to a CRD in the reference cell are part of a CRD in the query cell.

### CRD activity quantification

For CRD activity quantification, we applied a dimensionality reduction approach, i.e., we enumerated all ChIP-seq peaks per CRD, and took the mean of all single peak quantifications per individual to retrieve a single quantification value for each individual.

### Computing CRD-gene associations

We enumerated all the gene TSS within +/-1 Mb from the CRD boundaries and tested each one of these genes for association with the CRD activity. We retained the best adjusted nominal p-value for the number of genes being tested with 1000 permutations. To correct for the number of genes being tested, we used a false discovery rate (FDR) correction approach and declared phenotype-variant pairs at FDR 5% threshold as significant. These steps were carried out with QTLtools cis mode (https://qtltools.github.io/qtltools/) [40]. To discover multiple genes with independent effects on a given CRD, we used the conditional analysis approach implemented in QTLtools. Briefly, this approach is based on a forward-backward scan of the cis-window around the phenotypes to automatically find multiple independent gene-CRD associations, while controlling for a given FDR.

### CRD-gene associations specificity across cell types

We quantified CRD-gene cell type specific activities by considering the activity of the same exact set of peaks across the three cell types. The fraction of significant CRD-gene associations in common between the 2 cell types was then calculated by the number of significant CRD-gene associations in the query cell over the total number of significant CRD-gene associations of the reference cell (FDR 5%).

### CRD-Quantitative Trait Loci (CRD-QTL) mapping

For each CRD activity, we first enumerated all genetic variants within +/-1 Mb to the CRD boundaries and then tested each one of these variants for association with the phenotype and only retained the best hit (i.e., with the smallest nominal p-value). Secondly, we adjusted the best nominal p-value for the number of variants being tested by permutations. Specifically, we randomly shuffled the phenotype quantifications 1,000 times and retained the best association p-values for each permuted data set, which effectively gave 1,000 null p-values of associations. Third, to correct for the number of molecular phenotypes being tested throughout the genome we used a false discovery rate (FDR) correction approach and declared phenotype-variant pairs at FDR 5% threshold as significant. These steps were carried out with QTLtools cis mode (https://qtltools.github.io/qtltools/) [40]. To discover multiple QTLs with independent effects on a given molecular phenotype, we used the conditional analysis approach implemented in QTLtools. Briefly, this approach is based on a forwardbackward scan of the cis-window around the phenotypes to automatically learn the number of independent QTLs and to identify the most likely candidate variants, while controlling for a given FDR.

### CRD-Quantitative Trait Loci (CRD-QTL) specificity between cell types

To compare CRD-QTLs between cells, we considered the activity of the same exact set of peaks across the three cell types. We computed the significant CRD-QTL associations at 5% FDR is the reference cell type, and extracted these associations among all the CRD-QTL associations in the reference cell. From the adjusted p-values we computed the Pi1 estimate [25].

### TFBSs enrichment in CRD-QTLs

We incorporated knowledge from external databases such as Motifmap [26] and Remap [27], and extracted the 50 TFBSs that overlapped the most with ChIP-seq peaks located within CRDs. We computed enrichment for the significant CRD-QTLs within the 50 most represented TFBSs using QTLtools fenrich with 1000 permutations, and calculating subsequently Fisher’s test odds ratios at 1% FDR.

### PCHi-C support for molecular phenotype associations

CHiCAGO algorithms perform normalization and multiple testing specifically adapted to CHi-C experiments [29] and consider PCHi-C interactions significant if the interactions detected by CHiCAGO have a score superior or equal to 5. We integrate to our analysis the CHiCAGO score to PCHiC interaction matrices available for each cell type. We compute the number of associations supported by PCHi-C (with a CHiCAGO score ≥ 5) over the total number of associations, to obtain the fraction of associations supported by PCHi-C.

### CRD trans associations and Trans Regulatory Hubs

To map inter-chromosomal CRD-CRD associations, we performed association testing of all pairs of CRDs belonging to distinct autosomal chromosomes. We corrected for the number of tests by using the R/qvalue package and used a cutoff of 1% FDR threshold for downstream analyses. To call Trans Regulatory Hubs in a network of CRD-CRD associations, we used the sets of CRD-CRD associations at 1% FDR and detected communities using the R network package R/igraph with the greedy algorithm.

### Gene set enrichment analysis of genes associated to TRHs

To identify gene set enrichment in the TRHs, we extracted the genes associated to the CRDs involved in each TRH. We ran the GOrilla algorithm [1] for each TRH separately, on the 25 largest TRHs ranging from 8 to 3727 genes. We provided the algorithm with the target gene set (genes associated to the TRH) and a background gene set (all the genes expressed in each cell type). The GO enrichment terms analysis was performed using the default hypergeometric test after Benjamini–Hochberg false discovery rate correction for multiple testing set at 1% FDR, which led to 10 TRH remaining.

### Mapping trans eQTLs through histone Trans CRD network

To perform this analysis, we first assembled multiple layers of associations together in order to define three lists of gene-variant pairs to be tested for association in trans for each cell type:

scenario 1

1. aCRD-QTLs which links genetic variants and CRDs together (in cis),
2. CRD-CRD inter-chromosomal associations (in trans),
3. gene-CRD associations which links genes and CRDs together (in cis).

scenario 2

1. eQTLs which links genetic variants and genes together (in cis),
2. gene-CRD associations which links genes and CRDs together (in cis).
3. CRD-CRD inter-chromosomal associations (in trans),
4. gene-CRD associations which links genes and CRDs together (in cis).

Overlap of the discovered trans eQTLs with the significant trans eQTLs from the eQTLGen consortium [37] was performed using on pairs of variants in linkage disequilibrium (R2*>*0.5) as given by the API of the LDlink webtool (https://ldlink.nci.nih.gov) [41].

## 1 References

1. Maurano, Matthew T., et al. “Systematic localization of common disease-associated variation in regulatory DNA.” Science 337.6099 (2012): 1190–1195.

2. GTEx Consortium, Genetic effects on gene expression across human tissues. Nature 550, 204–213 (2017).

3. Hindorff, Lucia A., et al. “Potential etiologic and functional implications of genome-wide association loci for human diseases and traits.” Proceedings of the National Academy of Sciences 106.23 (2009): 9362–9367.

4. Chen, Lu, et al. “Genetic drivers of epigenetic and transcriptional variation in human immune cells.” Cell 167.5 (2016): 1398–1414.

5. Montgomery, Stephen B., and Emmanouil T. Dermitzakis. “From expression QTLs to personalized transcriptomics.” Nature Reviews Genetics 12.4 (2011): 277–282.

6. Lappalainen, Tuuli, et al. “Transcriptome and genome sequencing uncovers functional variation in humans.” Nature 501.7468 (2013): 506–511.

7. E. Lieberman-Aiden et al., Comprehensive mapping of longrange interactions reveals folding principles of the human genome. Science 326, 289–293 (2009).

8. J. Dekker, L. Mirny, The 3D genome as moderator of chromosomal communication. Cell 164, 1110–1121 (2016).

9. F. Spitz, E. E. Furlong, Transcription factors: From enhancer binding to developmental control. Nat. Rev. Genet. 13, 613–626 (2012).

10. E. S. Wong et al., Interplay of cis and trans mechanisms driving transcription factor binding and gene expression evolution. Nat. Commun. 8, 1092 (2017).

11. S. M. Waszak et al., Population variation and genetic control of modular chromatin architecture in humans. Cell 162, 1039–1050 (2015).

12. Delaneau, Olivier, et al. “Chromatin three-dimensional interactions mediate genetic effects on gene expression.” Science 364.6439 (2019).

13. Gate, R.E., Cheng, C.S., Aiden, A.P. et al. Genetic determinants of co-accessible chromatin regions in activated T cells across humans. Nat Genet 50, 1140–1150 (2018).

14. Wang, J., Zibetti, C., Shang, P. et al. ATAC-Seq analysis reveals a widespread decrease of chromatin accessibility in age-related macular degeneration. Nat Commun 9, 1364 (2018).

15. Bravo González-Blas, C., Minnoye, L., Papasokrati, D. et al. cisTopic: cis-regulatory topic modeling on single-cell ATAC-seq data. Nat Methods 16, 397–400 (2019).

16. Bendl, Jaroslav, et al., The three-dimensional landscape of chromatin accessibility in Alzheimer’s disease. BioRxiv (2021).

17. Chodavarapu, R. K., S. Feng,., M. Pellegrini. 2010. Relationship between nucleosome positioning and DNA methylation. Nature. 466:388–392.

18. Xu, Wanxue, et al. “Integrative analysis of DNA methylation and gene expression identified cervical cancer-specific diagnostic biomarkers.” Signal transduction and targeted therapy 4.1 (2019): 1–11.

19. Zhang, Ling, et al. “DNA methylation landscape reflects the spatial organization of chromatin in different cells.” Biophysical journal 113.7 (2017): 1395–1404.

20. Sahu, Biswajyoti, et al. “Sequence determinants of human gene regulatory elements.” Nature Genetics (2022): 1–12.

21. Martens, Joost HA, and Hendrik G. Stunnenberg. “BLUEPRINT: mapping human blood cell epigenomes.” Haematologica 98.10 (2013): 1487.

22. Javierre, Biola M., et al. “Lineage-specific genome architecture links enhancers and non-coding disease variants to target gene promoters.” Cell 167.5 (2016): 1369–1384.

23. Heintzman, Nathaniel D., et al. “Histone modifications at human enhancers reflect global cell-type-specific gene expression.” Nature 459.7243 (2009): 108–112.

24. Ribeiro, Diogo M., et al. “The molecular basis, genetic control and pleiotropic effects of local gene co-expression.” Nature communications 12.1 (2021): 1–13.

25. Storey, John D., and Robert Tibshirani. “Statistical significance for genomewide studies.” Proceedings of the National Academy of Sciences 100.16 (2003): 9440–9445.

26. Daily, Kenneth, et al. “MotifMap: integrative genome-wide maps of regulatory motif sites for model species.” BMC bioinformatics 12.1 (2011): 1–13.

27. Chèneby, Jeanne, et al. “ReMap 2018: an updated atlas of regulatory regions from an integrative analysis of DNA-binding ChIP-seq experiments.” Nucleic acids research 46.D1 (2018): D267–D275.

28. Uhlen, Mathias, et al. “Towards a knowledge-based human protein atlas.” Nature biotechnology 28.12 (2010): 1248–1250.

29. Cairns, Jonathan, et al. “CHiCAGO: robust detection of DNA looping interactions in Capture Hi-C data.” Genome biology 17.1 (2016): 1–17.

30. Freire-Pritchett, P., Ray-Jones, H., Della Rosa, M. et al. Detecting chromosomal interactions in Capture Hi-C data with CHiCAGO and companion tools. Nat Protoc 16, 4144–4176 (2021).

31. Grundberg, Elin, et al. “Mapping cis-and trans-regulatory effects across multiple tissues in twins.” Nature genetics 44.10 (2012): 1084–1089.

32. Csardi, Gabor, and Tamas Nepusz. “The igraph software package for complex network research.” InterJournal, complex systems 1695.5 (2006): 1–9.

33. Eden, Eran, et al. “GOrilla: a tool for discovery and visualization of enriched GO terms in ranked gene lists.” BMC bioinformatics 10.1 (2009): 1–7

34. Supek, Fran, et al. “REVIGO summarizes and visualizes long lists of gene ontology terms.” PloS one 6.7 (2011): e21800.

35. Cooke, Michael P., et al. “Regulation of T cell receptor signaling by a src family protein-tyrosine kinase (p59fyn).” Cell 65.2 (1991): 281–291

36. Rosales, Carlos. “Neutrophil: a cell with many roles in inflammation or several cell types?.” Frontiers in physiology 9 (2018): 113.

37. Võsa, Urmo, et al. “Unraveling the polygenic architecture of complex traits using blood eQTL metaanalysis.” BioRxiv (2018): 447367.

38. Wingett, Steven, et al. “HiCUP: pipeline for mapping and processing Hi-C data.” F1000Research 4 (2015).

39. Heinz, Sven, et al. “Simple combinations of lineage-determining transcription factors prime cisregulatory elements required for macrophage and B cell identities.” Molecular cell 38.4 (2010): 576–589.

40. Delaneau, Olivier, et al. “A complete tool set for molecular QTL discovery and analysis.” Nature communications 8.1 (2017): 1–7.

41. Alexander TA, Machiela MJ. LDpop: an interactive online tool to calculate and visualize geographic LD patterns. BMC Bioinformatics. 2020 Jan 10.

