## Supplementary Figures for "Genetic variation in correlated regulatory region of Immunity"

Supplementary Table 1A: CRDs and CRD associations (CRD-QTLs, CRD-genes, CRD-CRD) discovered in Monocytes, Neutrophils and T-cells

| Type of CRD | Cell Type | Sample Size | CRDs discovered | Sample Size | CRDs discovered |
|---|---|---|---|---|---|
| hCRD | Neutrophils | 94 | 6,831 | 165 | 7,666 |
| | Monocytes | 94 | 7,660 | 160 | 9,287 |
| | T-cells | 94 | 5,480 | 94 | 5,480 |
| | LCLs | 94 | 10,497 | 317 | 12,583 |
| mCRD | Neutrophils | 94 | 3,828 | 197 | 6,112 |
| | Monocytes | 94 | 3,877 | 196 | 6,053 |
| | T-cells | 94 | 4,275 | 132 | 5,701 |

| Type of CRD | Type of QTL | Description | Range | Number of CRDs tested in these associations | | | Number of discoveries at 5% FDR | | |
|---|---|---|---|---|---|---|---|---|---|
| | | | | Monocytes | Neutrophils | T-cells | Monocytes | Neutrophils | T-cells |
| hCRD | CRD-QTL | Genetic variant associated with CRD activity (5% FDR) | Cis | 9,287 CRDs | 7,666 CRDs | 5,480 CRDs | 7,050 | 4,854 | 1,616 |
| | CRD-gene | Gene associated with CRD activity (5% FDR) | Cis | | | | 6,755 | 6,300 | 2,239 |
| | CRD-CRD | CRD-CRD associations (1% FDR) | Trans | | | | 84,690 | 159,422 | 116,658 |
| | TRH | cluster of trans CRD-CRD associations | Trans | | | | 308 | 107 | 31 |
| mCRD | CRD-QTL | Genetic variant associated with CRD activity (5% FDR) | Cis | 6,053 CRDs | 6,112 CRDs | 5,701 CRDs | 4,325 | 4,363 | 3,786 |
| | CRD-gene | Gene associated with CRD activity (5% FDR) | Cis | | | | 2,300 | 2,027 | 1,858 |
| | CRD-CRD | CRD-CRD associations (1% FDR) | Trans | | | | 11,719 | 12,525 | 14,022 |
| | TRH | cluster of trans CRD-CRD associations | Trans | | | | 230 | 234 | 262 |

S1B

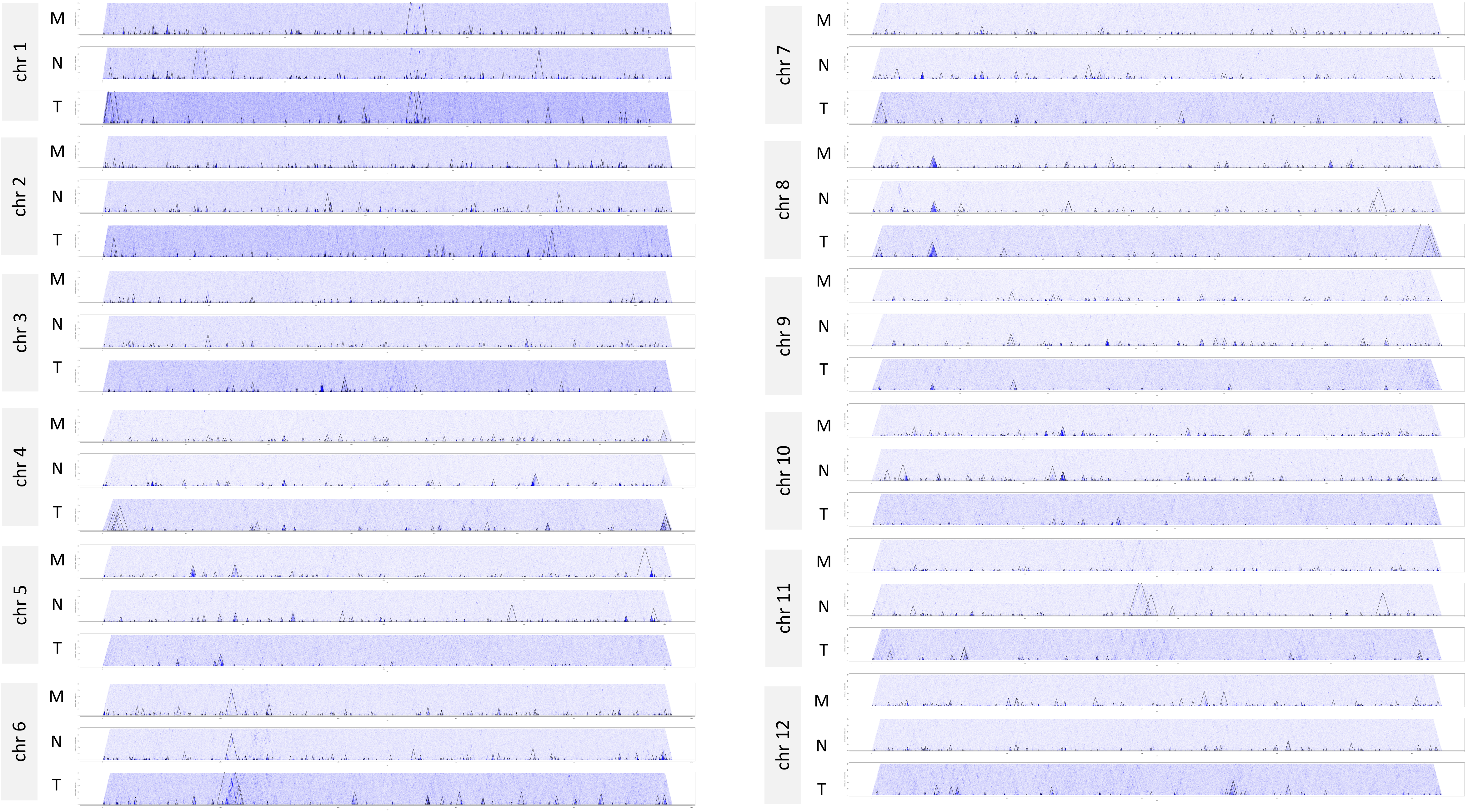
chr 1
M
N
T
chr 2
M
N
T
chr 3
M
N
T
chr 4
M
N
T
chr 5
M
N
T
chr 6
M
N
T
chr 7
M
N
T
chr 8
M
N
T
chr 9
M
N
T
chr 10
M
N
T
chr 11
M
N
T
chr 12
M
N
T

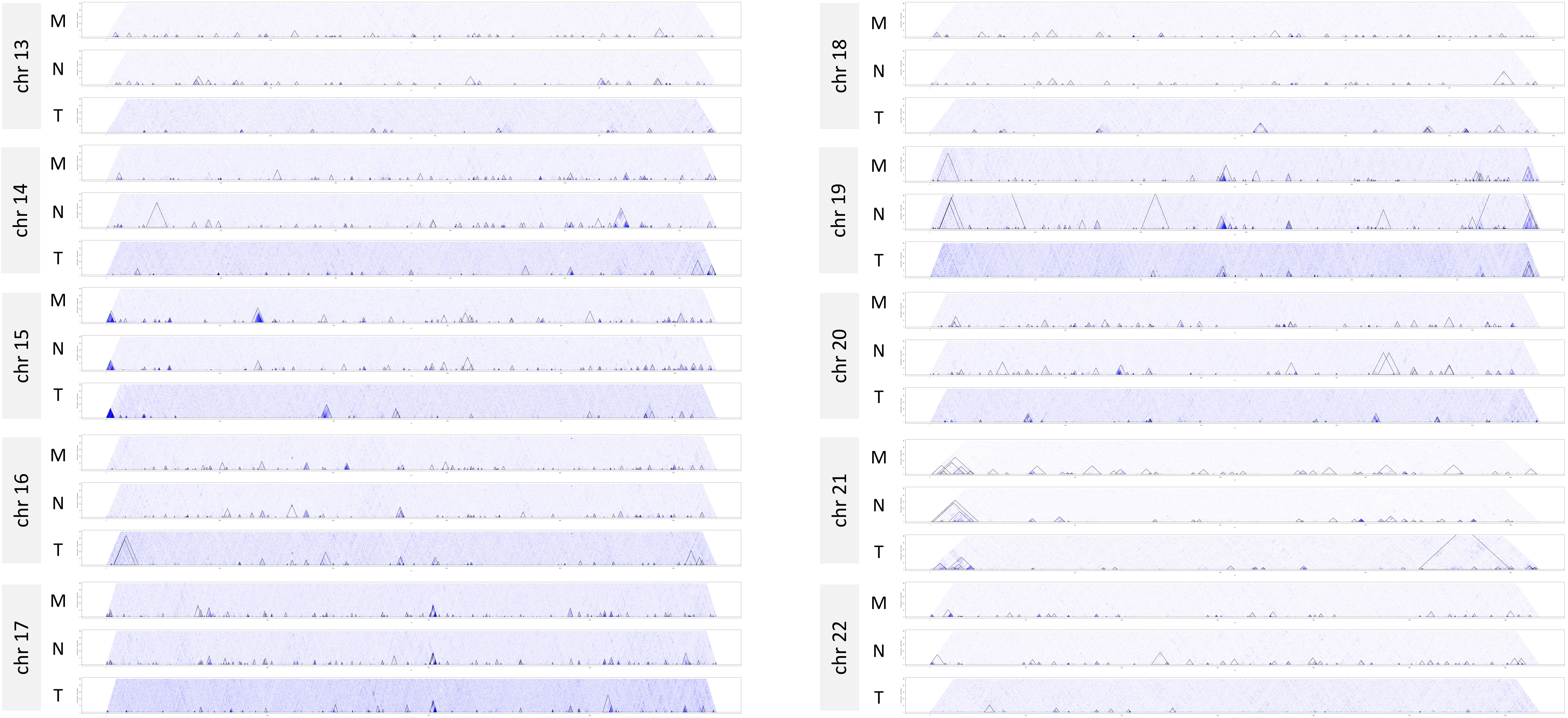
chr 13
M
N
T
chr 14
M
N
T
chr 15
M
N
T
chr 16
M
N
T
chr 17
M
N
T
chr 18
M
N
T
chr 19
M
N
T
chr 20
M
N
T
chr 21
M
N
T
chr 22
M
N
T

S1C

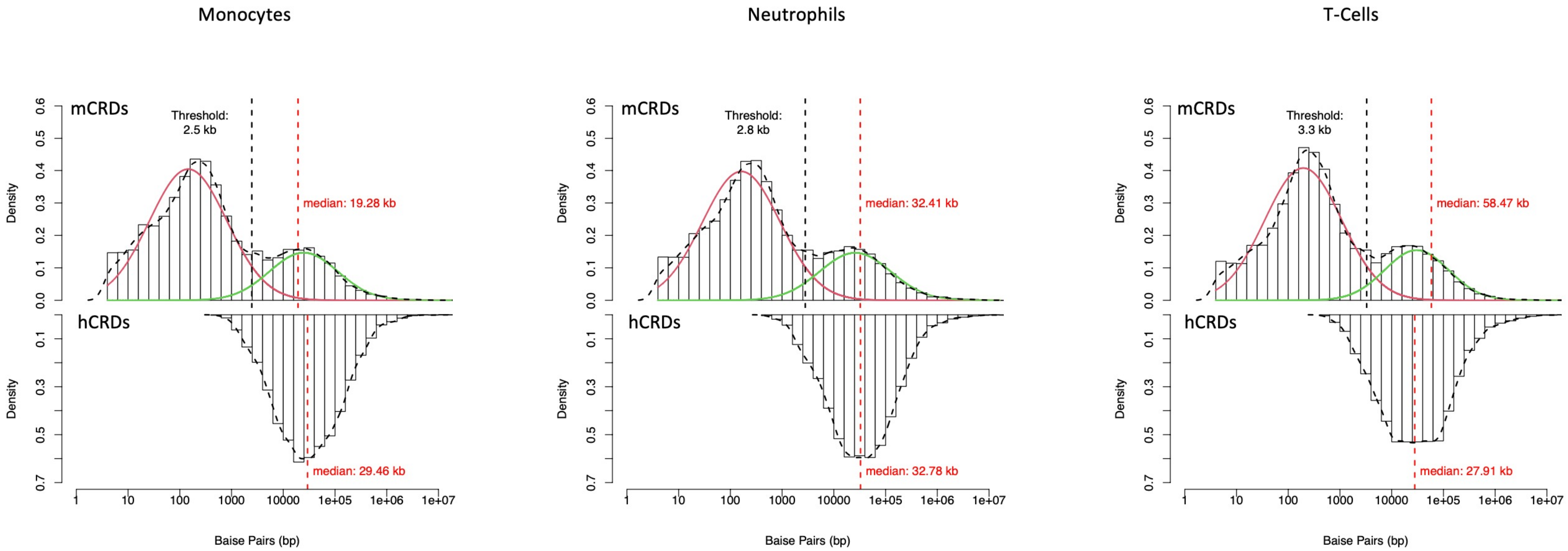

Monocytes
Neutrophils
T-Cells
mCRDs
hCRDs
Threshold:
2.5 kb
Threshold:
2.8 kb
Threshold:
3.3 kb
median: 19.28 kb
median: 29.46 kb
median: 32.41 kb
median: 32.78 kb
median: 58.47 kb
median: 27.91 kb
Density
Baise Pairs (bp)

S1D

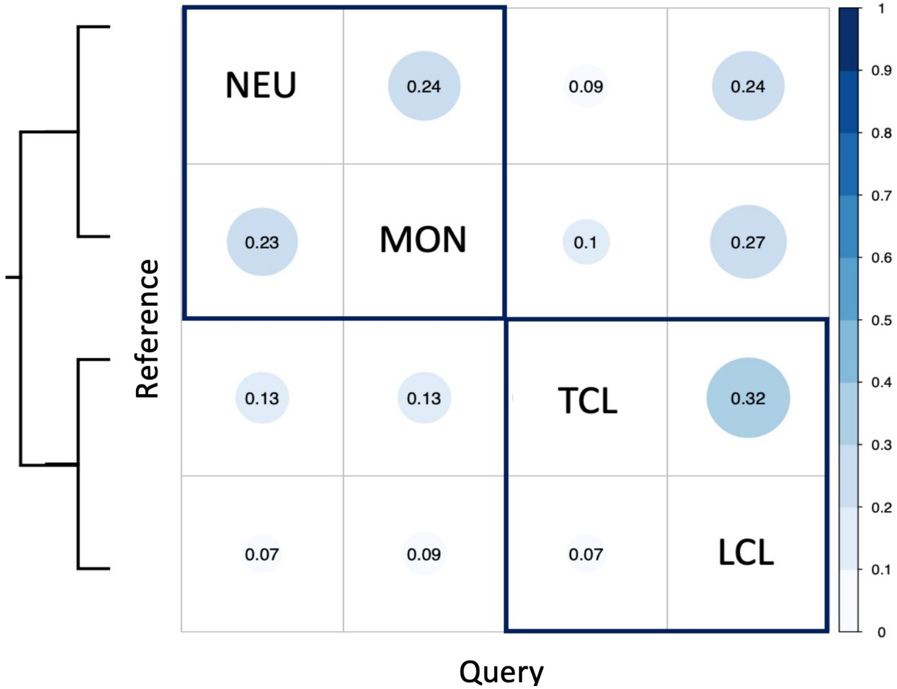

S1E

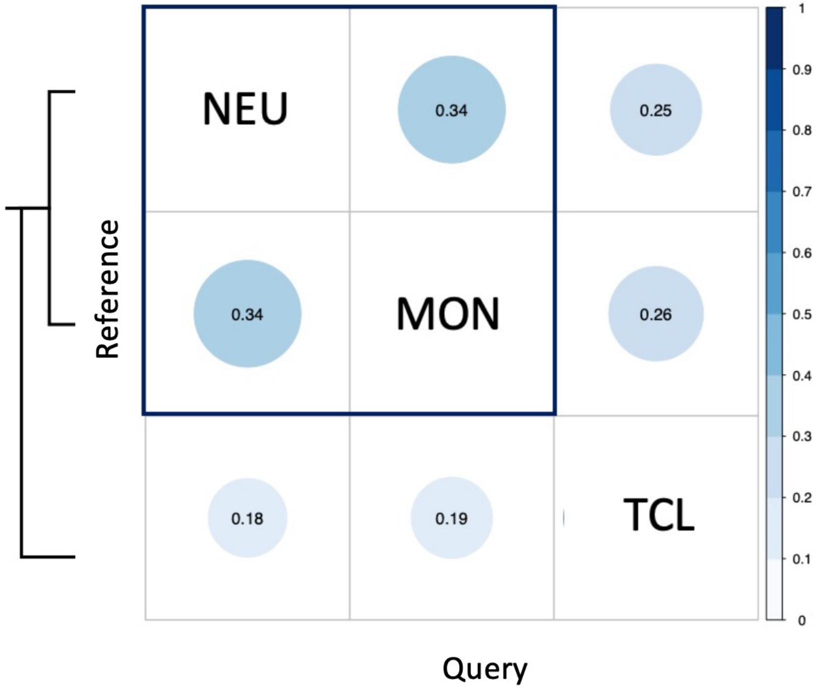

S1F

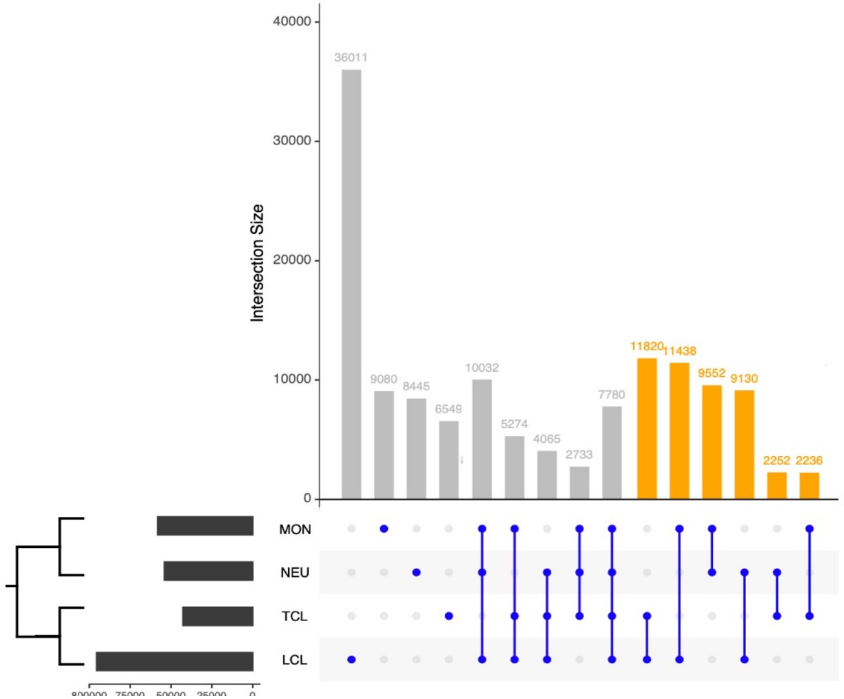

S1G

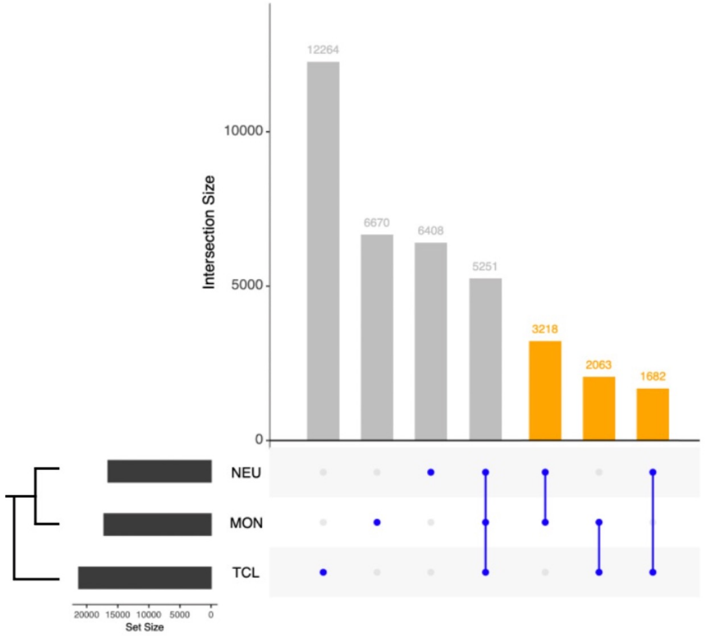

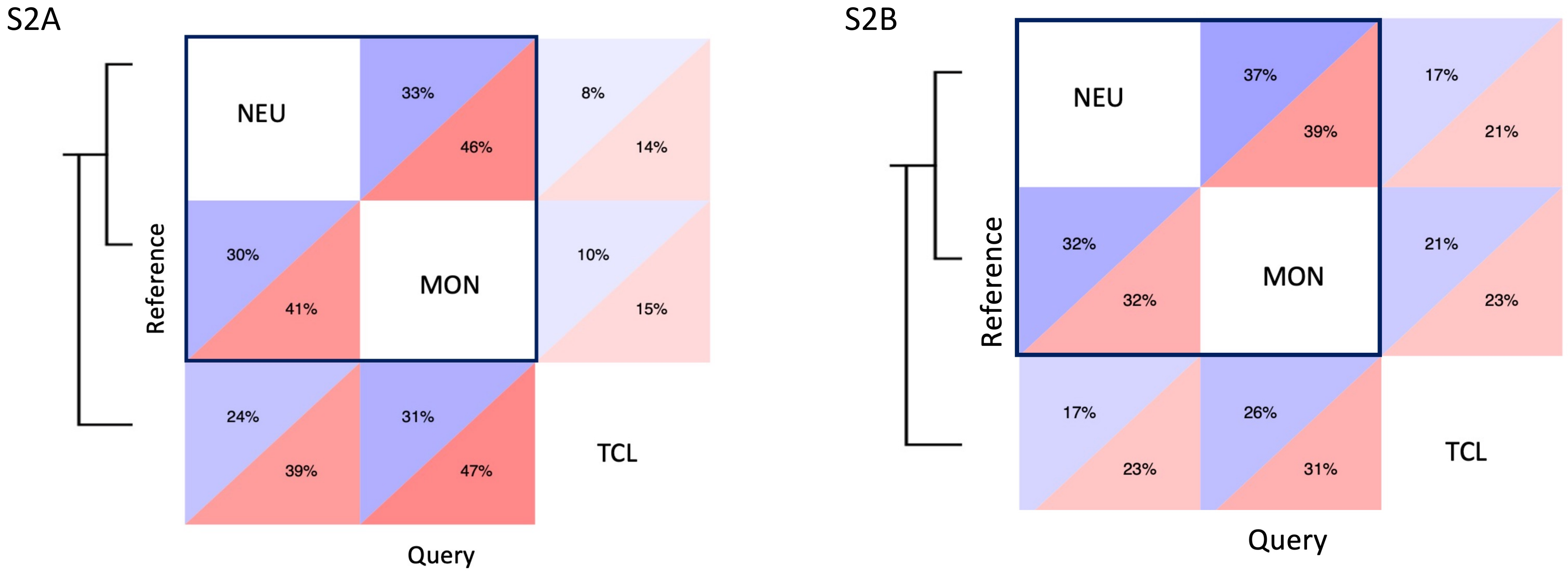
S2A
NEU
33%
46%
8%
14%
30%
41%
MON
10%
15%
24%
39%
31%
47%
TCL
Reference
Query
S2B
NEU
37%
39%
17%
21%
32%
32%
MON
21%
23%
17%
23%
26%
31%
TCL
Reference
Query

S2C

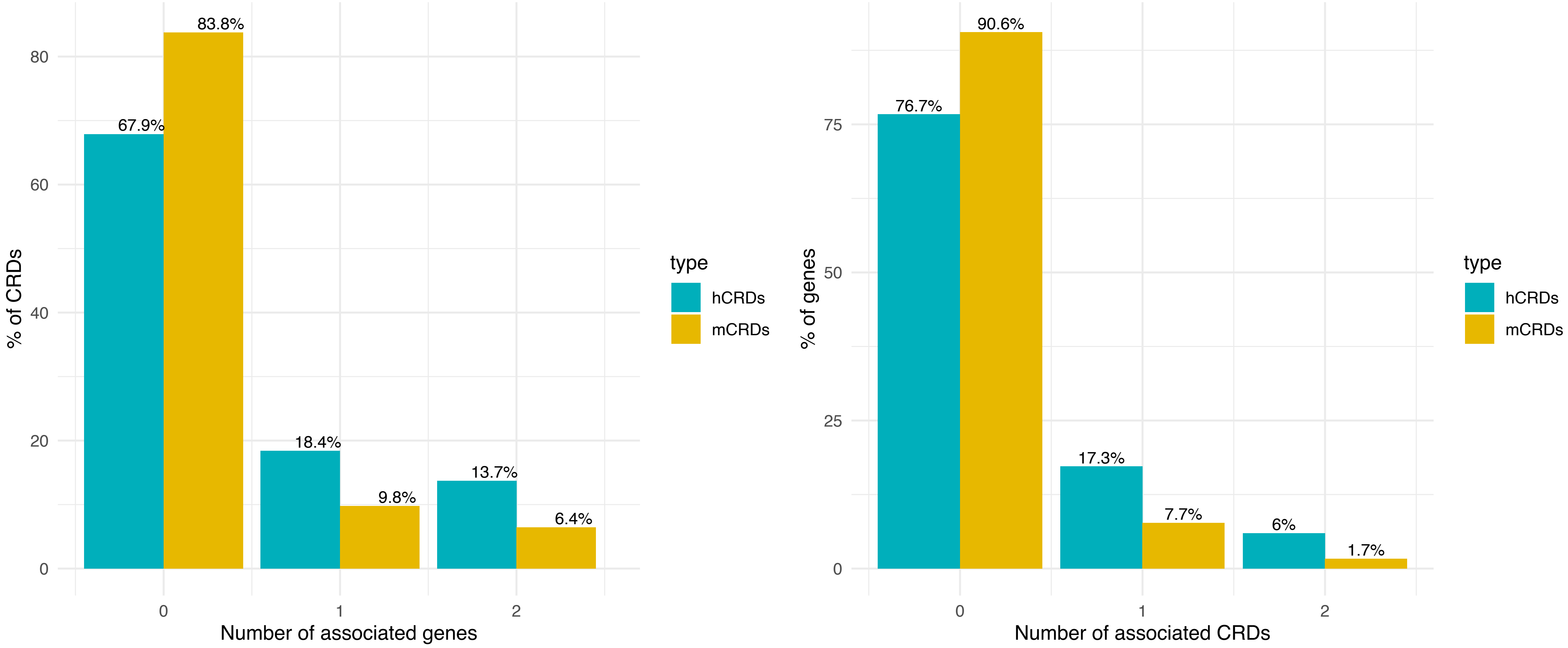

S2D

| | Co-Expressed Gene Pairs | |
|---|---|---|
| Cell Type | Tested | FDR<0.01 |
| Neutrophils | 4,663,276 | 29,940 |
| Monocytes | 6,663,972 | 46,146 |
| T-cells | 5,872,671 | 13,737 |

S2E

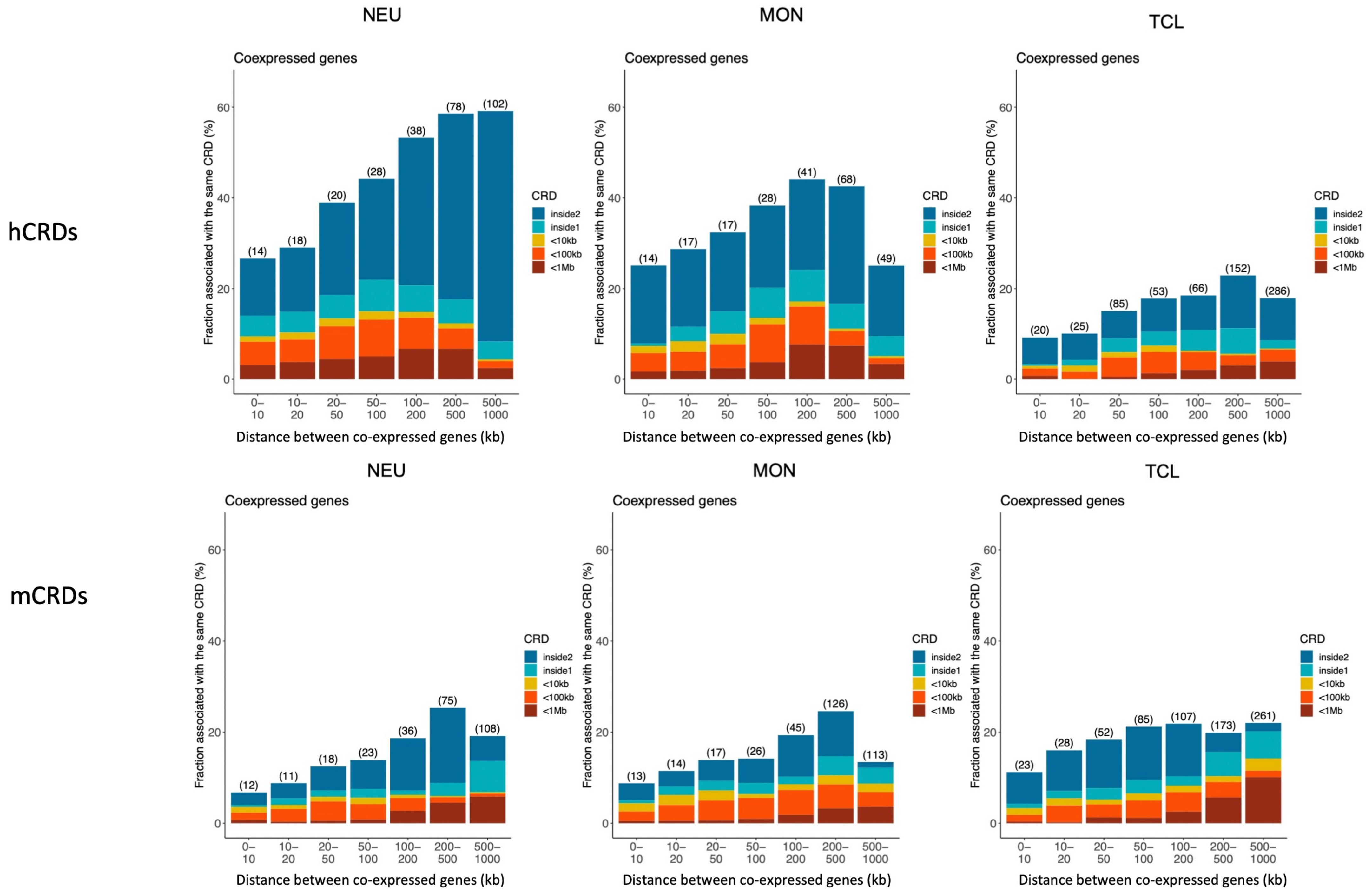
hCRDs
mCRDs
NEU
MON
TCL
Coexpressed genes
Fraction associated with the same CRD (%)
Distance between co-expressed genes (kb)
0–10
10–20
20–50
50–100
100–200
200–500
500–1000
CRD
inside2
inside1
<10kb
<100kb
<1Mb
(14) (18) (20) (28) (38) (78) (102)
(14) (17) (17) (28) (41) (68) (49)
(20) (25) (85) (53) (66) (152) (286)
(12) (11) (18) (23) (36) (75) (108)
(13) (14) (17) (26) (45) (126) (113)
(23) (28) (52) (85) (107) (173) (261)

S2F

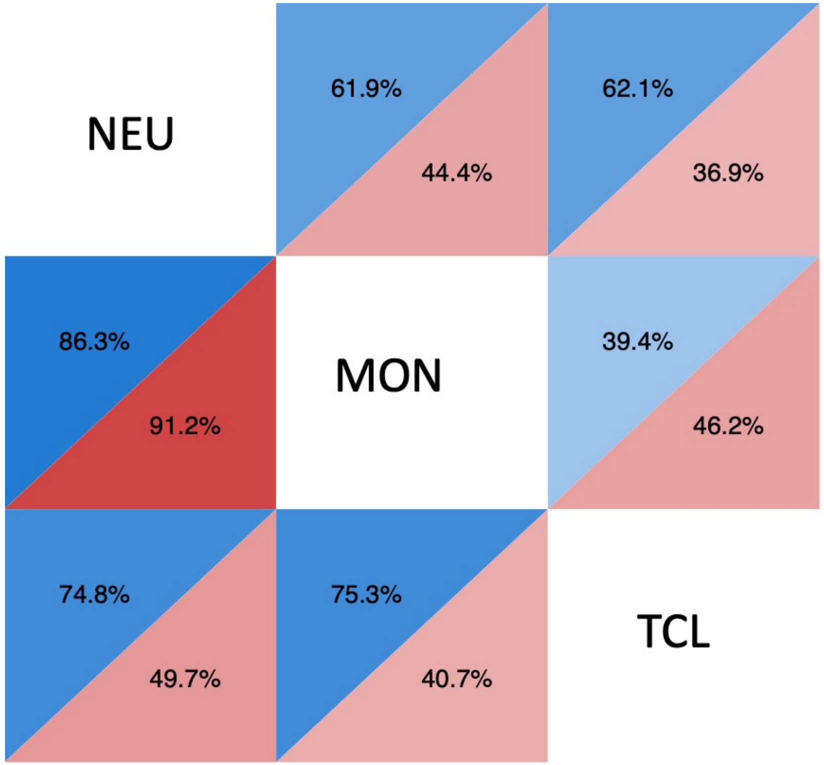

S2G

| data_type | cell | TFBS | pval | oddsratio |
|---|---|---|---|---|
| hist | mono | SPI1 | 8,84E-90 | 3,48 |
| hist | neut | SPI1 | 3,11E-47 | 3,08 |
| hist | tcell | TCF4 | 1,49E-07 | 3,04 |
| methyl | mono | ETS2 | 4,12E-02 | 3,00 |
| hist | tcell | SPI1 | 1,25E-15 | 2,93 |
| hist | mono | STAT1 | 1,48E-61 | 2,82 |
| hist | mono | TCF4 | 1,91E-21 | 2,77 |
| methyl | neut | SPI1 | 2,19E-29 | 2,75 |
| methyl | tcell | TCF4 | 8,57E-11 | 2,61 |
| methyl | mono | SPI1 | 1,01E-24 | 2,52 |
| methyl | mono | TCF4 | 1,80E-11 | 2,49 |
| methyl | neut | TCF4 | 1,25E-10 | 2,46 |
| hist | tcell | BCL11A | 2,16E-07 | 2,43 |
| hist | neut | STAT1 | 7,40E-30 | 2,43 |
| methyl | mono | MED12 | 7,73E-07 | 2,40 |
| hist | mono | BCL11A | 6,57E-24 | 2,39 |
| methyl | tcell | SPI1 | 3,14E-18 | 2,38 |
| methyl | neut | BCL11A | 1,04E-15 | 2,34 |
| hist | mono | STAT3 | 8,71E-56 | 2,33 |
| hist | tcell | STAT1 | 1,11E-09 | 2,29 |
| methyl | mono | PU.1 | 3,06E-03 | 2,26 |
| methyl | tcell | BCL11A | 7,96E-13 | 2,26 |
| methyl | mono | BCL11A | 1,62E-14 | 2,23 |
| hist | neut | TCF4 | 1,26E-08 | 2,19 |
| methyl | neut | GABP | 1,24E-18 | 2,16 |
| methyl | neut | STAT1 | 1,50E-19 | 2,14 |
| methyl | neut | SOX4 | 4,44E-06 | 2,13 |
| methyl | mono | SOX4 | 4,22E-06 | 2,11 |
| hist | tcell | GABP | 2,02E-06 | 2,10 |
| methyl | tcell | MED12 | 3,01E-04 | 2,09 |
| hist | neut | BCL11A | 1,79E-11 | 2,06 |
| methyl | tcell | STAT1 | 3,91E-15 | 2,06 |
| methyl | mono | GABP | 3,71E-16 | 2,03 |
| hist | neut | STAT3 | 2,47E-25 | 2,01 |
| hist | mono | GABP | 1,10E-18 | 2,00 |

S2H

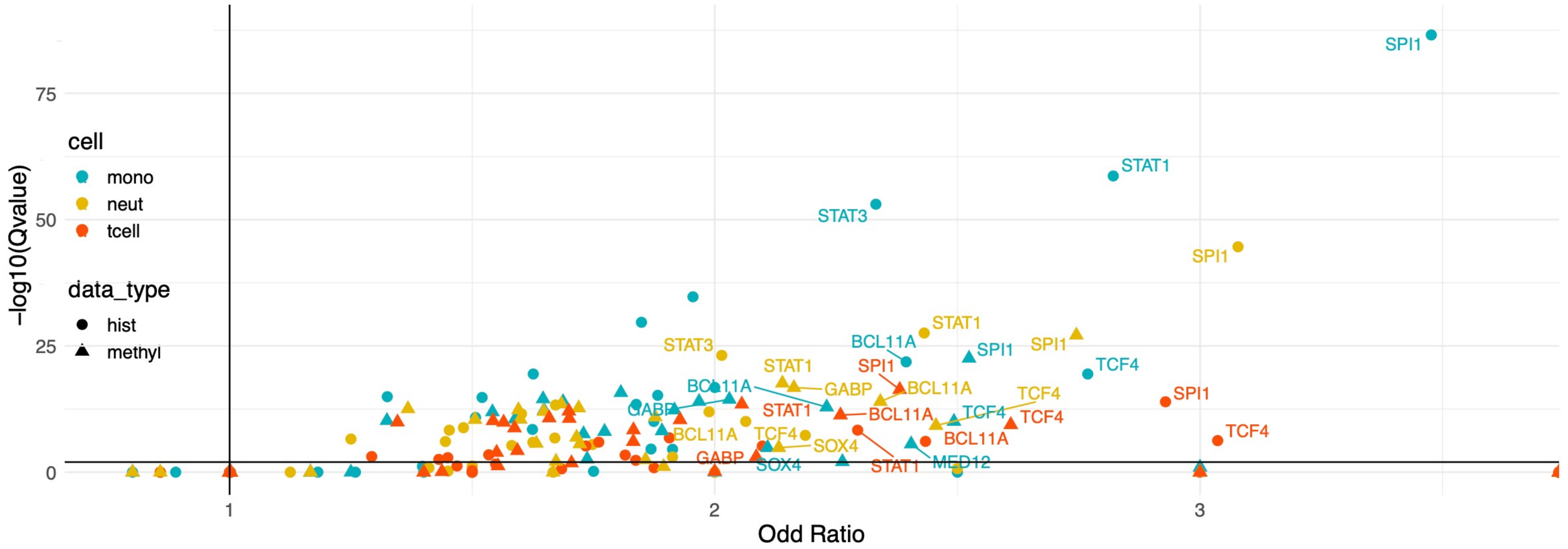

S3A S3B S3C

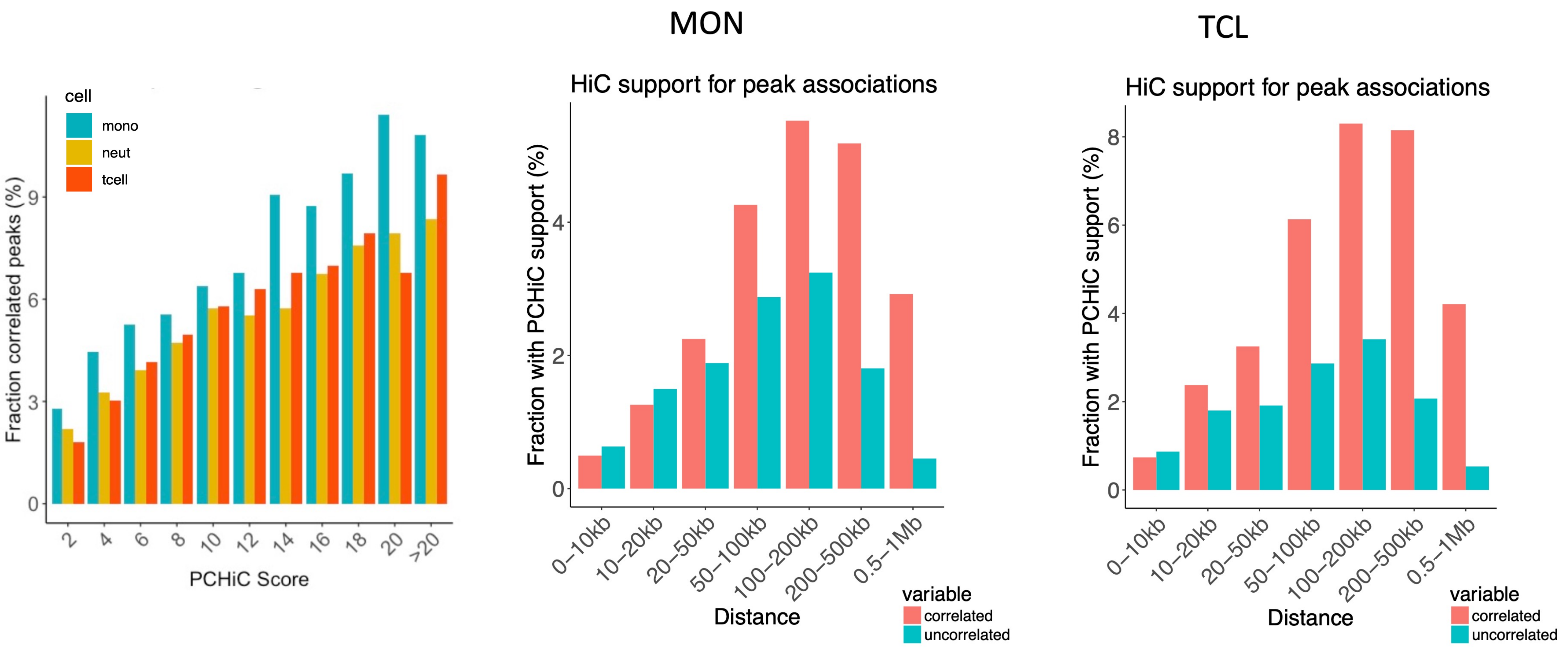
cell
mono
neut
tcell
Fraction correlated peaks (%)
0
3
6
9
2
4
6
8
10
12
14
16
18
20
>20
PCHiC Score
MON
HiC support for peak associations
Fraction with PCHiC support (%)
0
2
4
0–10kb
10–20kb
20–50kb
50–100kb
100–200kb
200–500kb
0.5–1Mb
Distance
variable
correlated
uncorrelated
TCL
HiC support for peak associations
Fraction with PCHiC support (%)
0
2
4
6
8
0–10kb
10–20kb
20–50kb
50–100kb
100–200kb
200–500kb
0.5–1Mb
Distance
variable
correlated
uncorrelated

S3D

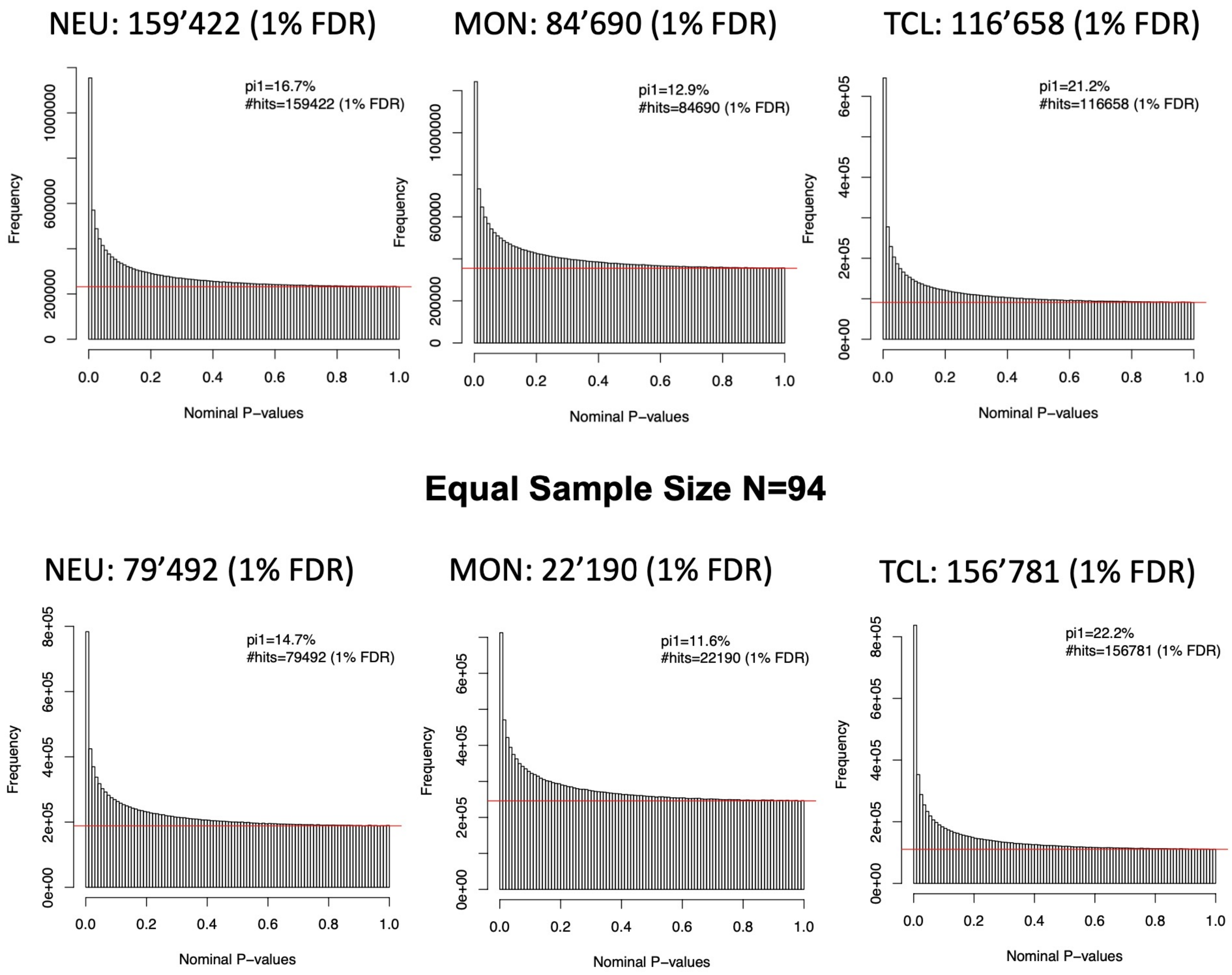
NEU: 159’422 (1% FDR)
MON: 84’690 (1% FDR)
TCL: 116’658 (1% FDR)
pi1=16.7%
#hits=159422 (1% FDR)
pi1=12.9%
#hits=84690 (1% FDR)
pi1=21.2%
#hits=116658 (1% FDR)
Frequency
Nominal P–values
Equal Sample Size N=94
NEU: 79’492 (1% FDR)
MON: 22’190 (1% FDR)
TCL: 156’781 (1% FDR)
pi1=14.7%
#hits=79492 (1% FDR)
pi1=11.6%
#hits=22190 (1% FDR)
pi1=22.2%
#hits=156781 (1% FDR)
Frequency
Nominal P–values

S3E

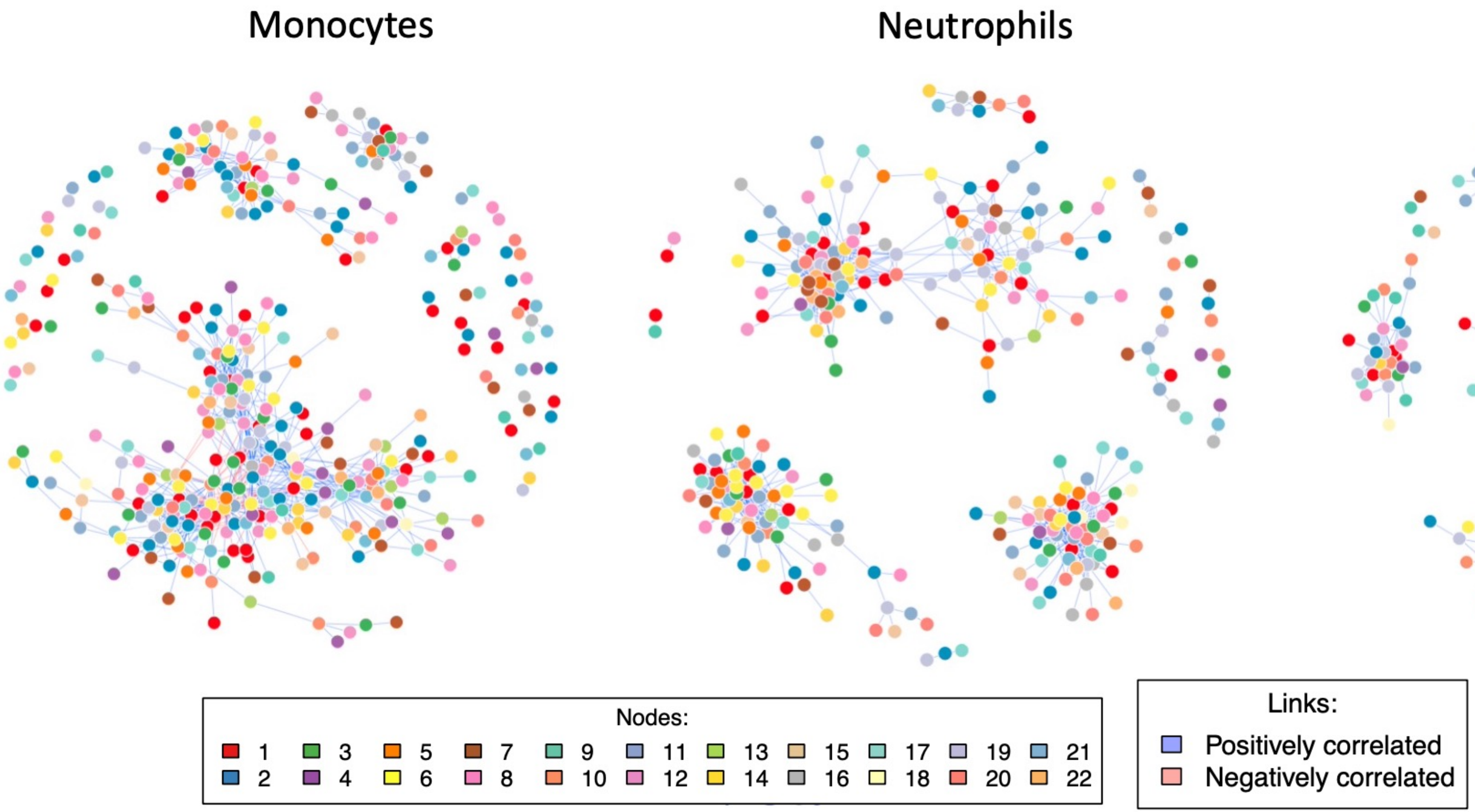
Monocytes
Neutrophils
T-cells
Nodes:
Links:
Positively correlated
Negatively correlated

S3F

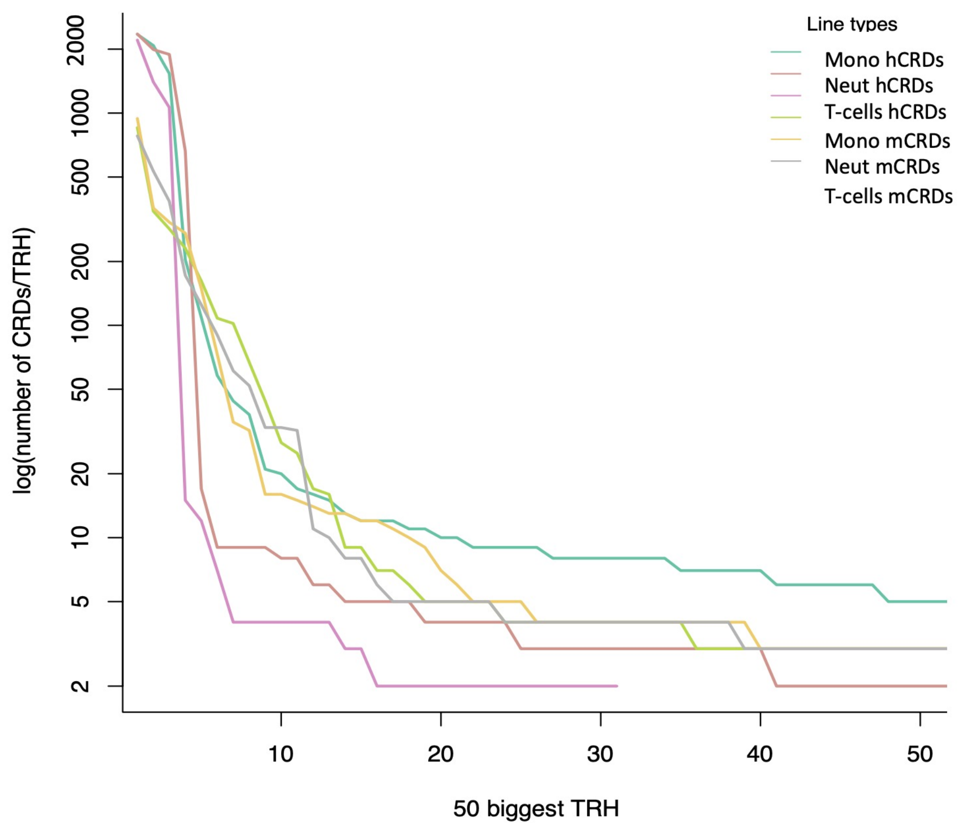
log(number of CRDs/TRH)
2000
1000
500
200
100
50
20
10
5
2
10
20
30
40
50
50 biggest TRH
Line types
Mono hCRDs
Neut hCRDs
T-cells hCRDs
Mono mCRDs
Neut mCRDs
T-cells mCRDs

S3G

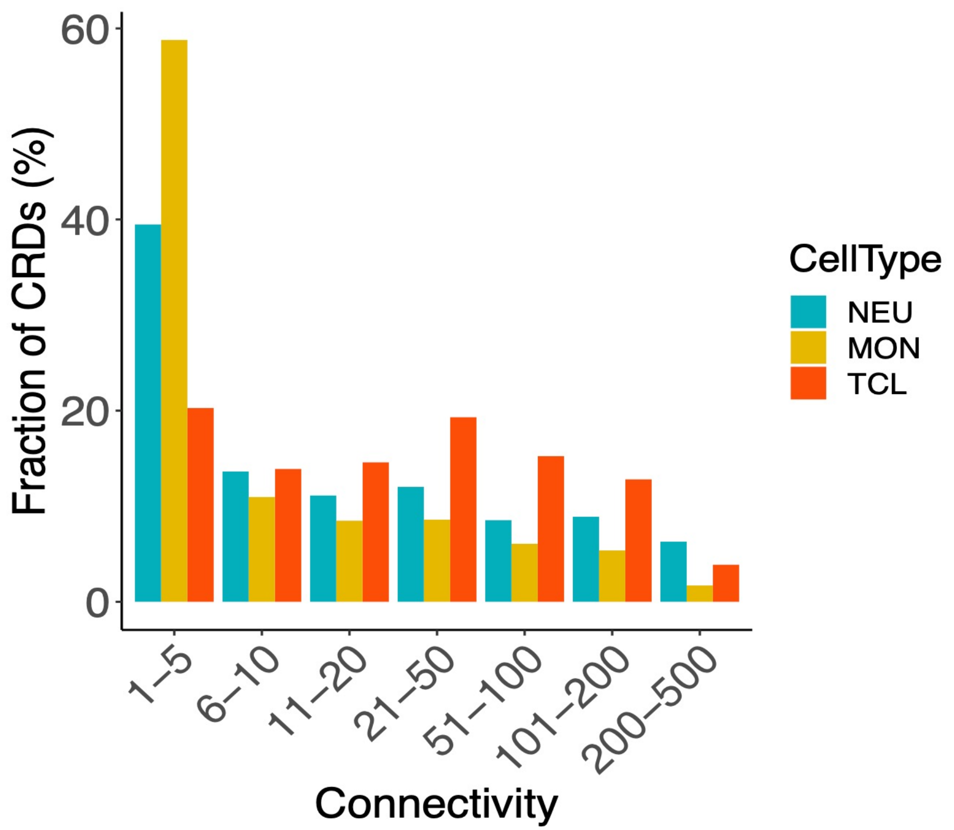
Fraction of CRDs (%)
60
40
20
0
1–5
6–10
11–20
21–50
51–100
101–200
200–500
Connectivity
CellType
NEU
MON
TCL

S3H

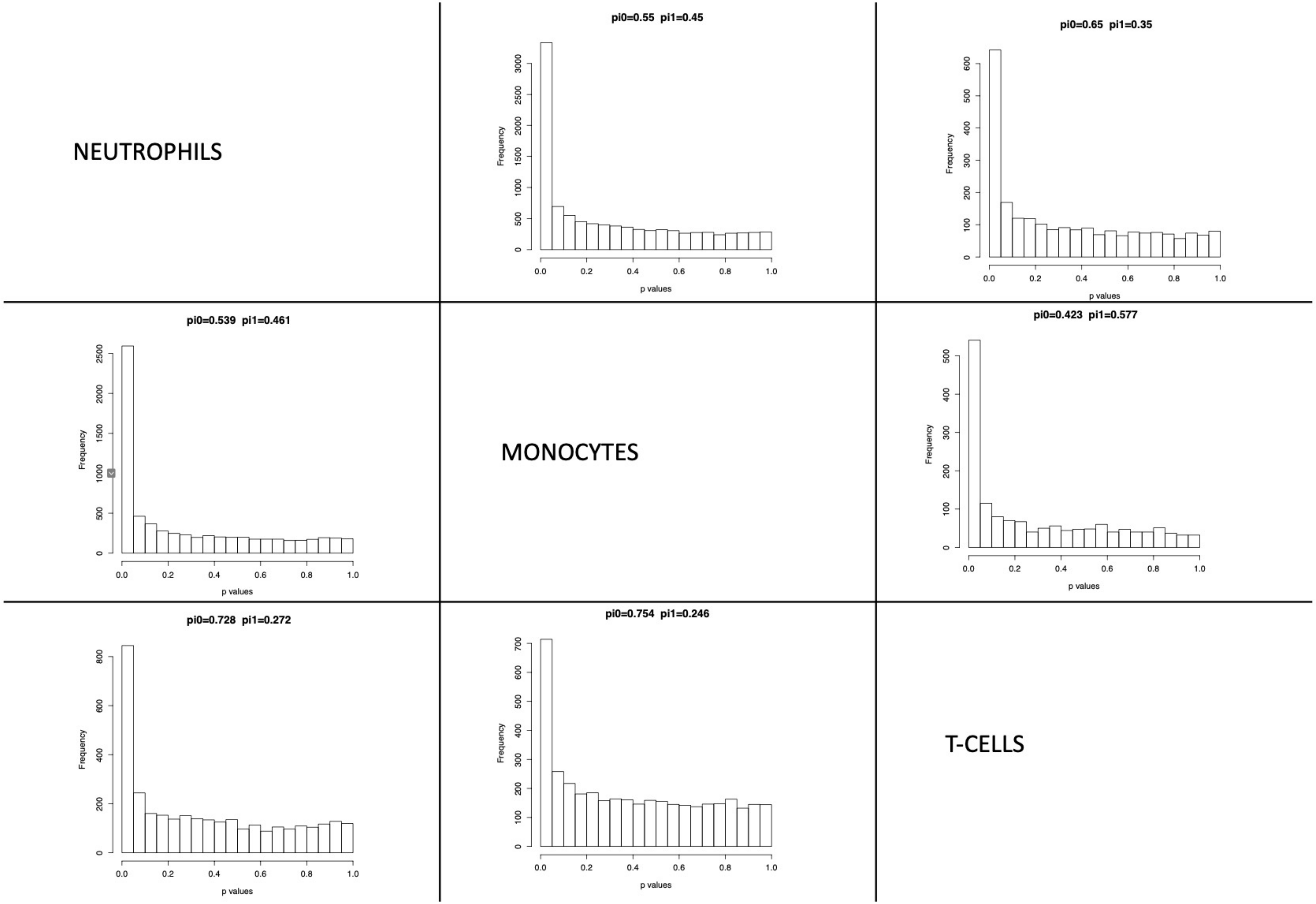
NEUTROPHILS
MONOCYTES
T-CELLS
pi0=0.55 pi1=0.45
pi0=0.65 pi1=0.35
pi0=0.539 pi1=0.461
pi0=0.423 pi1=0.577
pi0=0.728 pi1=0.272
pi0=0.754 pi1=0.246
Frequency
p values

S3I

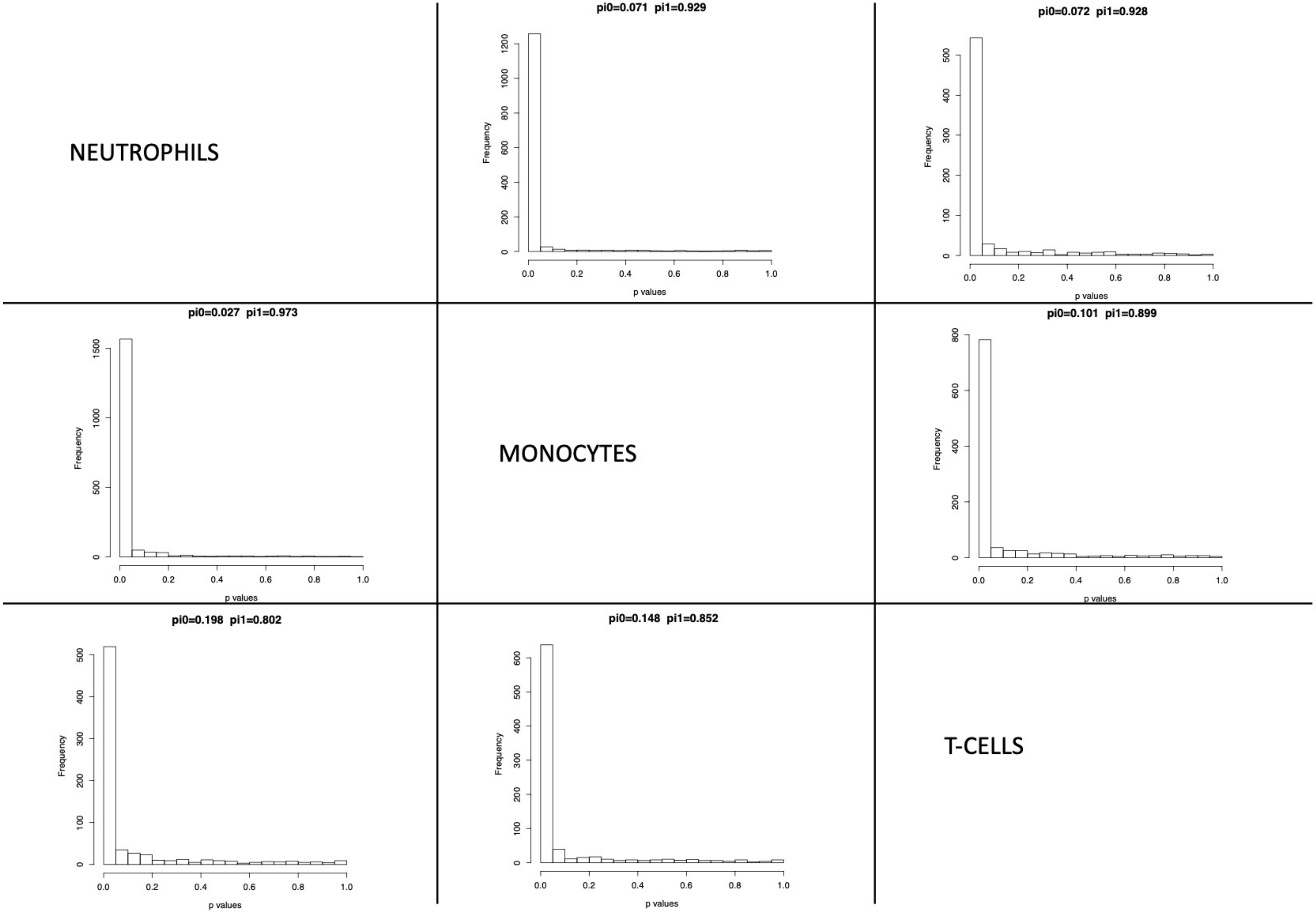
NEUTROPHILS
pi0=0.071 pi1=0.929
pi0=0.072 pi1=0.928
pi0=0.027 pi1=0.973
MONOCYTES
pi0=0.101 pi1=0.899
pi0=0.198 pi1=0.802
pi0=0.148 pi1=0.852
T-CELLS
Frequency
p values

S3J

| Cell | GOTerm | description | Enrichment | Qvalue |
|---|---|---|---|---|
| T-cells | GO:0045959 | negative regulation of complement activation, classical pathway | 30.9 | 0.00723 |
| T-cells | GO:0030450 | regulation of complement activation, classical pathway | 30.9 | 0.00734 |
| T-cells | GO:0045619 | regulation of lymphocyte differentiation | 3.98 | 0.000738 |
| T-cells | GO:0002822 | regulation of adaptive immune response based on somatic recombination of immune receptors built from immunoglobulin superfamily domains | 3.83 | 0.00745 |
| T-cells | GO:0002819 | regulation of adaptive immune response | 3.64 | 0.00691 |
| T-cells | GO:0002250 | adaptive immune response | 3.49 | 0.00929 |
| T-cells | GO:0030098 | lymphocyte differentiation | 3.41 | 0.00745 |
| T-cells | GO:1902105 | regulation of leukocyte differentiation | 3.01 | 0.005 |
| T-cells | GO:0002521 | leukocyte differentiation | 2.9 | 0.00963 |
| T-cells | GO:0045087 | innate immune response | 2.7 | 0.00256 |
| T-cells | GO:0002683 | negative regulation of immune system process | 2.62 | 0.00266 |
| T-cells | GO:0050778 | positive regulation of immune response | 2.6 | 5.11e-05 |
| T-cells | GO:0002684 | positive regulation of immune system process | 2.55 | 3.89e-07 |
| T-cells | GO:0050776 | regulation of immune response | 2.53 | 5.55e-07 |
| T-cells | GO:0002253 | activation of immune response | 2.5 | 0.00477 |
| T-cells | GO:0006955 | immune response | 2.42 | 9.82e-05 |
| T-cells | GO:0002682 | regulation of immune system process | 2.34 | 7.79e-09 |
| Neutrophils | GO:0002682 | regulation of immune system process | 1.43 | 0.0026 |
| Neutrophils | GO:0002376 | immune system process | 1.35 | 0.00314 |

S3K

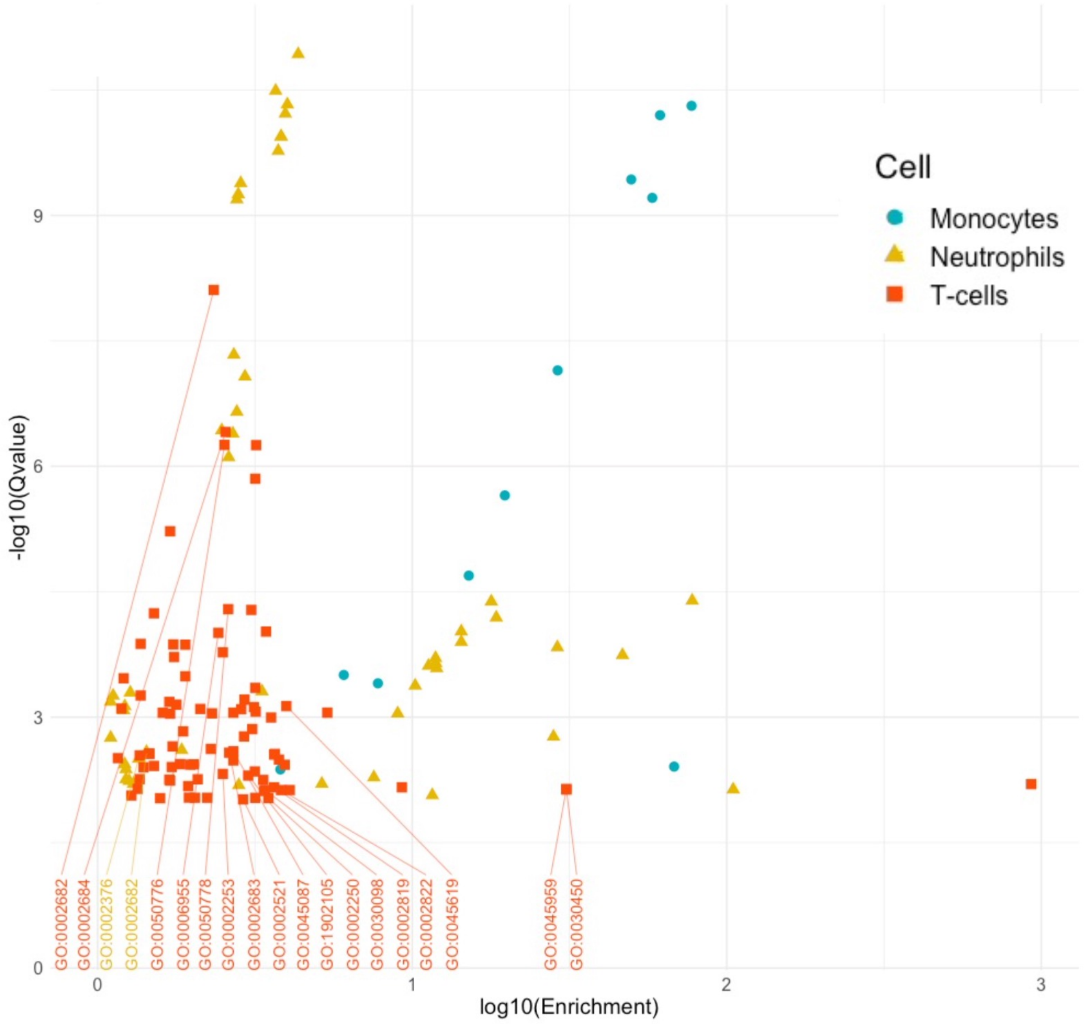

S3L

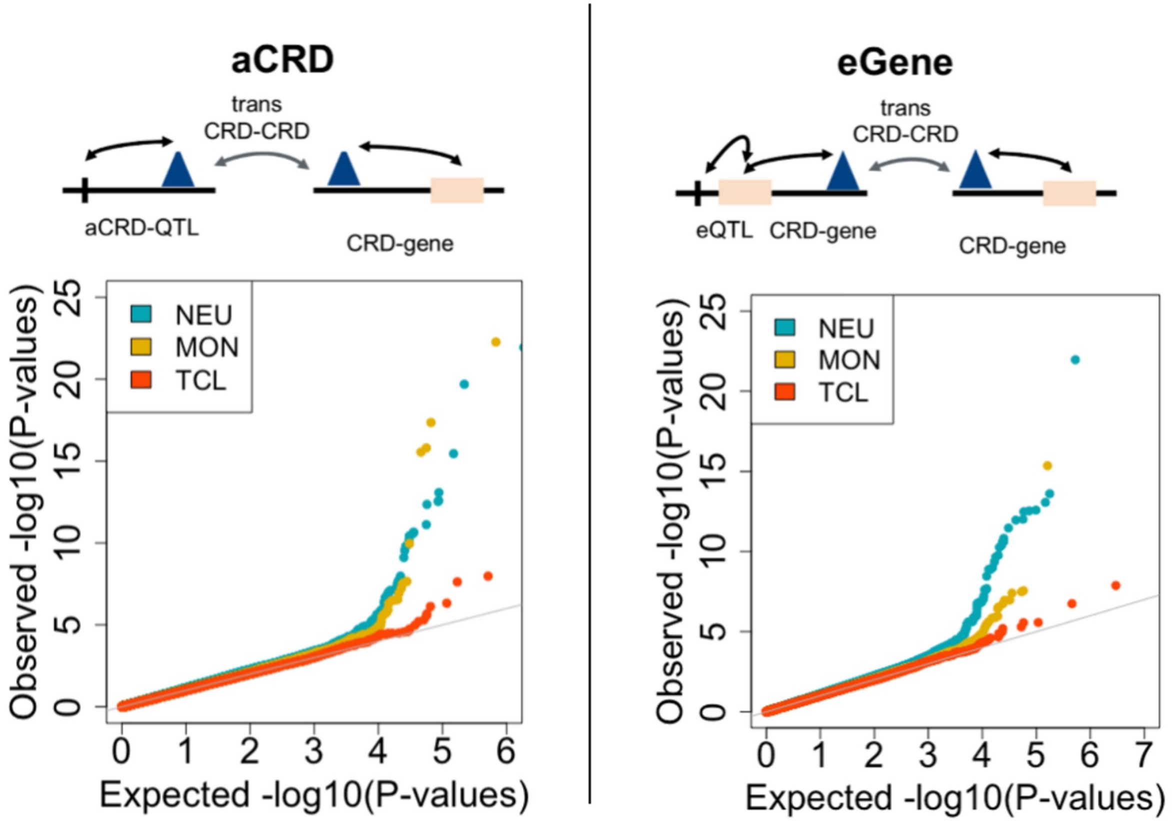

S3M

| | **Trans eQTL mapping** | | | |
|---|---|---|---|---|
| | **Scenario 1** | | **Scenario 2** | |
| Cell Type | Tested | FDR<0.05 | Tested | FDR<0.05 |
| Neutrophils | 265,055 | 55 | 305,359 | 62 |
| Monocytes | 176,392 | 16 | 164,610 | 65 |
| T-cells | 45,767 | 5 | 53,420 | 4 |

S3N

| | **Overlap with eQTL Gen trans-eQTLs (R2>0.5)** | | | |
|---|---|---|---|---|
| | Unique hits FDR<0.05 | Overlap eQTLGen associations | Genes with overlap | Variants with overlap |
| Neutrophils | 117 | 40 | 24 | 4 |
| Monocytes | 81 | 1 | 1 | 1 |
| T-cells | 9 | 1 | 1 | 1 |
