## Supplementary Legends for "Genetic variation in correlated regulatory region of Immunity"

### Figure Legends

#### Regulatory interaction dynamics across immune cell types

Supplementary 1A:

CRDs and CRD associations (CRD-QTLs, CRD-genes, CRD-CRD) discovered in Monocytes, Neutrophils, T-cells and LCLs.

Supplementary 1B:

hCRDs across neutrophils, monocytes, T-cells for 3 cell types.

Supplementary 1C:

Size distribution of mCRDs (top) and mCRDs (bottom) in Neutrophils, Monocytes and T-cells. The median is represented in red and the modelling of mCRD distribution with a mixture gaussian model of small domains (0.2 to a few kb) and large domains (a few kb to 1-2Mb) is displayed in red and green.The threshold corresponding to the 0.95 percentile is plotted in black.

Supplementary 1D,E:

Tissue sharing of hCRDs (D) and mCRDs (E) across neutrophils, monocytes and T-cells for an identical sample size of 94 samples. A CRD is shared between two cell types more than 50% of the peaks of the reference CRD are present in a CRD from the query tissue. The fraction of sharing is shown for every combination of tissues.

Supplementary 1F,G:

Overlap across cell types for chromatin peaks (F) and CpG marks (E) belonging to hCRDs

#### CRDs are under genetic control

Supplementary 2A,B:

hCRD-gene(A) and mCRD-gene(B) association sharing (blue) at 5% FDR. A CRD-gene association is shared between two cell types if the reference CRD and gene are also associated in the query cell type. The fraction of sharing is shown for every combination of tissues. Fraction of genes shared between cell-types (red). We considered the genes involved in the hCRD-gene associations (5% FDR)

Supplementary 2C:

Fraction of CRDs or genes as a function of the number of genes or CRDs they are associated with, respectively for hCRDs, mCRDs in neurophils, monocytes and T-cells.

Supplementary Table 2D:

Cis co-expressed gene pairs (i.e. genes whose expression is correlated among individuals) (FDR 1\%) respectively for neutrophils, monocytes and T-cells.

Supplementary 2E:

Percentages of pairs of co-expressed genes (5%FDR) that associate with the same hCRD(top) / mCRD (bottom) as a function of distance between genes. Gene pairs odds ratios of belonging to the same CRD while being co-expressed are shown in parentheses. Each bar is stratified by the mean distance between pair of genes and their associated CRDs. The position of the genes according to the CRD is color coded on the figure.

Supplementary 2F:

$\pi_1$ estimate of hCRD-QTL (blue) and mCRD-QTL (red) sharing between cell-types

Supplementary 2G:

Table summarizing the enrichment in TFBS for significant CRD-QTLs, with odd ratio>2 and p-value <0.05.

Supplementary 2H:

Enrichment in the 50 most represented TFs for significant CRD-QTLs. Labels present for TFBS with an odd ratio > 2 and p-value>$10^{-4}$

### CRD structure and connectivity reflect functional 3D chromatin organization

Supplementary 3A:

Percentages of significant association at 1% FDR between pairs of peaks located on the same chromosome as a function of PCHi-C signal (CHiCAGO score).

Supplementary 3B:

Fraction of monocytes chromatin peak pairs on the same chromosome supported by PCHi-C data (CHiCAGO score >5) at significantly associated (pink) and non-associated (blue) pairs of chromatin peaks within bins of increasing distance between peaks.

Supplementary 3C:

Fraction of T-cells chromatin peak pairs on the same chromosome supported by PCHi-C data (CHiCAGO score >5) at significantly associated (pink) and non-associated (blue) pairs of chromatin peaks within bins of increasing distance between peaks.

### Trans CRD networks highlight cell-type specific biological functions in immune cells

Supplementary 3D:

Distribution of nominal p-values for CRD-CRD trans associations in neutrophils, monocytes and T cells. The number of significant associations at FDR=1% is given above each histogram. On the bottom, same figures for a similar sample size (n=94) across cell-types.

Supplementary 3E:

Network representation of the 1,000 strongest associations between histone CRDs on distinct chromosomes for neutrophils, monocytes and T cells. Each network node is a CRD and each edge is a significant association between two CRDs. Nodes are colored by chromosome number.

Supplementary 3F:

Distribution of histone and methyl TRH sizes for a maximum of 50 TRHs across cell types (top) and methyl TRH sizes (bottom)

Supplementary 3G:

Connectivity of hCRDs in CRD-CRD trans networks across cell types.

Supplementary 3H,I :

Patterns of trans hCRD-hCRD (H) and trans mCRD-mCRD (I) association sharing were also estimated by extracting the significant trans CRD associations identified in one cell-type and then tested for replication in another cell type

Supplementary 3J:

Top 20 Gene Ontology terms linked to immune response, from 2 TRHs: one in neutrophils and one in T-cells.

Supplementary 3K:

Gene Set Enrichment Analysis of genes belonging in a TRH. Enrichment of GO Term as a function of q- value for 3 cell types.

#### Mapping trans eQTLs through histone Trans CRD networks unravel new trans eQTLs

Supplementary 3L:

Schematic of the different data integration strategies to link variants to genes on different chromosomes and QQplots of trans-eQTL nominal P-values for neutrophils, monocytes and T cells.

Supplementary 3M:

Significant trans-eQTL (FDR 5%) found for each cell-type.

Supplementary 3N:

Overlap between trans eGenes (aCRD and eGene) across the three immune cell types, with eQTLGen trans-eQTLs.
